# Social tolerance as strategy to resolve group conflicts: an experimental study on tug-of-war competition of burying beetles

**DOI:** 10.1101/2022.12.08.519695

**Authors:** Long Ma, Wenxia Wang, Denise Roffel, Marco van der Velde, Jan Komdeur

## Abstract

In animal groups with dominance hierarchies, there often occurs a tug-of-war competition over resources and reproduction between dominants and subordinates, because neither is able to fully control the other. Consequently, individuals may mitigate within-group conflict, either by fighting others or by signalling their willingness to tolerate others. Nevertheless, how such a tolerance interaction evolves remains unclear. Here, we addressed this knowledge gap and tested the tug-of-war competition hypothesis, by investigating whether subordinates pay to stay in the group by helping dominants (pay-to-stay), and whether dominants pay costs by living with subordinates in the group (pay-from-staying). We used the burying beetles, *Nicrophorus vespilloides*, which compete with intra- and inter-specifics for valuable carcasses that are needed for reproduction. Multiple conspecifics can reproduce together through communal breeding, thereby enhancing benefits in terms of reproduction and resource defence against competitors. In communal associations, larger individuals are often dominant in carcass use and reproduction, whereas subordinates have restricted access to the carcass. Our findings show that cooperative subordinates paid costs by helping dominant breeders in carcass preparation in order to be tolerated (i.e. increased access towards the carcass) by dominant breeders, but subordinates did not increase their reproductive success by helping. Such tolerance was eliminated by a high interspecific competition with blowfly maggots. Our results also show that dominant males, but not dominant females, benefitted more from the presence of subordinates, partly due to a sex difference in the compensation strategy of dominants. Overall, our study demonstrates that a social tolerance occurring in situations with a tug-of-war competition could be a common strategy to resolve conflicts in animal societies.

## Introduction

Conspecific individuals often form temporary or permanent groups due to resource- and/or reproduction-sharing, by which they may gain fitness pay-offs in the short- or long-term (reviewed by ***Beekman et al., 2003***; ***Neukom et al., 2011***; ***Ratnieks et al., 2006***; ***West and Ghoul, 2019***). However, intra-group conflict of interests over resources and reproduction can still be manifested underneath the peaceful surface of social interactions, and this could incur different costs for each group member in terms of reproduction and other fitness traits (***Lukas and Clutton-Brock, 2012***; ***Whitehouse and Lubin, 1999***). Even though the resolution of such conflict promotes cooperation in animal societies and has attracted much attention (***Cameron et al., 2009***; ***Conradt and Roper, 2009***; ***Duboscq et al., 2014***; ***Ridley and Raihani, 2008***; ***Shen et al., 2017***), whether and how group members make a compromise for the allocation of resources and reproduction, thereby maximizing individual fitness benefits, as well as group productivity, remains less studied (***Aureli et al., 2002***; ***Cant and Young, 2013***; ***Conradt and Roper, 2009***; ***Young and Clutton-Brock, 2006***).

In some social groups, such as cooperative and communal breeding, high-ranked individuals take advantages in resource monopolization and likely become dominant in reproductive allocation between individuals (***Liu, Chan, et al., 2020***; ***Young and Clutton-Brock, 2006***). In contrast, other members (e.g. subordinates) may gain direct and/or indirect benefits from staying within groups, such as access to resources and mates, reproduction (***Heg et al., 2009***; ***Reeve and Keller, 2001***; ***Zöttl, Fischer, et al., 2013***) and low predation risk (***Groenewoud et al., 2016***; ***Krams et al., 2010***). However, dominant members often pay costs from living with subordinates because of a high intensity of intraspecific competition over food and other resources (***Hamilton, 2013***; ***Hamilton and Taborsky, 2005***). Subsequently, subordinate members may be coerced to pay costs by helping dominants, for instance, by building a communal nest (***Chen et al., 2020***; Leighton and Meiden, 2016), or by providing alloparental care (***Grinsted and Field, 2018***; ***Zöttl, Heg, et al., 2013***). Such helping behaviour may occur to gain kin-selected benefits, wherein individuals help their relatives reproduce (***Hamilton, 1964***; ***Komdeur and Hatchwell, 1999***; ***Nam et al., 2010***; ***Zöttl, Heg, et al., 2013***). In contrast, the ‘pay-to-stay’ theory offers an alternative explanation for the helping behaviour of individuals when the relatedness between actors and recipients is low (***Bergmüller et al., 2007***; ***Gaston, 1978***; ***Hellmann and Hamilton, 2018***; ***Koenig and Walters, 2011***; ***Kokko et al., 2002***). In particular, subordinates pay by helping in order to remain in the group and to obtain delayed direct benefits, such as the inheritance of territory (***Heg et al., 2009***; ***Kingma, 2017***) and the acquisition of breeding experience (***Jennions and Macdonald, 1994***; ***Richardson et al., 2007***). Also, they might be paying to gain direct benefits such as opportunities to breed concurrently with dominants (that is, paying-to-breed), or to become new breeding partners for ‘widowed’ individuals that have lost their breeding partner(***Balshine-Earn et al., 1998***; ***Bergmüller and Taborsky, 2005***). In many situations, the payment by subordinates seems not to fully compensate for the costs that they impose on dominants, for instance, when a breeding society consists of non-relatives, or resources are scarce and limited (***Eggert and Sakaluk, 2000***; ***Hamilton and Taborsky, 2005***; ***Liu, Chan, et al., 2020***; ***Young and Clutton-Brock, 2006***). Despite subordinates’ contribution towards group defence, their reproduction (e.g. more offspring produced) may negatively affect dominants’ reproduction on a limited breeding resource that can only support a certain number of offspring (***Eggert and Sakaluk, 2000***; ***Cant and Young, 2013***). It also means that, irrespective of whether or not subordinates help as a form of payment, dominants suffer costs from living with subordinates in the group, such as lower reproductive success (***Bergmüller and Taborsky, 2005***; ***Gilchrist et al., 2004***; ***Heg et al., 2009***). Consequently, a cooperative and peaceful relationship between dominants and subordinates has been suggested to avoid the escalation of within-group conflicts, thereby profiting each group member (***Boyd et al., 2003***; ***Clutton-Brock and Parker, 1995***).

In essence, the extent to which reproduction is partitioned between group members is expected to influence the payoffs of group breeding, as well as the level of cooperation in groups (***Hamilton et al., 2005***; ***Holekamp and Strauss, 2016***; ***Ratnieks et al., 2006***). The reproductive skew theory provides a prediction of the reproductive partitioning among group members, which generally consists of two main theoretical models (***Clutton-Brock, 1998***; ***Hamilton, 2013***; ***Johnstone, 2000***; ***Reeve and Keller, 2001***). The ‘transactional’ model assumes a complete control of one group member over the total group reproduction, which accounts for a stable ‘social agreement’ among individuals who have differences in reproductive incentives and some fitness traits (***Hamilton, 2013***; ***Reeve et al., 1998***). For example, in some highly eusocial Hymenoptera societies, worker reproduction is rare and sterile workers help fertile queens to rear their offspring (***Reeve and Keller, 2001***). Alternatively, the ‘tug-of-war’ model highlights a more common scenario, in which neither of each group member have full control over the reproduction of others, and an increased allocation to one individual’s reproduction may decrease the reproduction of others, and even overall group productivity (***Hamilton, 2013***; ***Reeve et al., 1998***; ***Shen and Reeve, 2010***). The extent to which each individual reproduces in one shared brood accounts for a lasting struggle over resources and reproduction, which is largely dependent on their competitive ability (***Bradley et al., 2005***; ***Hamilton, 2013***; ***Langer et al., 2004***; ***Shen and Reeve, 2010***). Moreover, the inability of dominants to control subordinate’s reproduction correlates with ecological and intrinsic conditions (***Creel and Waser, 1991***; ***Monnin and Peeters, 1999***). For instance, there is no or moderate skew in reproduction when resources are abundant or interspecific competition pressure is low, whereas in the reverse case reproduction is highly skewed (***Creel and Waser, 1997***; ***Eggert and Müller, 1992***; ***Langer et al., 2004***). Hitherto, although the ‘tug-of-war’ phenomena are widespread across animal species, the theoretical ‘tug-of-war’ model is little studied empirically (***Bradley et al., 2005***; ***Cant, 2000***; ***Gilchrist, 2006***; ***Gilchrist et al., 2004***; ***Hamilton, 2013***). This study aims to unravel the mechanism behind the tug-of-war competition model and how individuals adjust the level of cooperation and conflict in groups, using the burying beetle, *Nicrophorus vespilloides*. These beetles always breed and take care of their offspring in pairs, and can communally breed on single resources, which is an ideal model for scrutinizing the tug-of-war prediction in animal societies (***Eggert and Sakaluk, 2000***; ***Liu, Chan, et al., 2020***; ***Müller et al., 2007***; ***Shen and Reeve, 2010***).

Burying beetles search for and prepare small vertebrate carcasses as food and breeding resources. Female beetles always breed alone, and they often cooperatively bury a carcass and breed in pairs. Then, they raise offspring and provide extended care on the buried carcass until offspring develop to independence (***Eggert and Müller, 2011***; ***Müller et al., 2007***; ***Scott, 1998***; ***Scott and Panaitof, 2004***; ***Smiseth et al., 2005***). Multiple individuals of conspecifics can often prepare a carcass together and then breed communally in a group (***Eggert and Sakaluk, 2000***; ***Komdeur et al., 2013***; ***Müller et al., 2007***; ***Richardson and Smiseth, 2020***). For each individual, joining in a group is proposed to enhance the synergetic capacity of obtaining resources and defending them against competitors (e.g. blowfly; ***Chen et al., 2020***; ***Liu, Chan, et al., 2020***; ***Sun et al., 2014***). Additional group members might potentially be beneficial because of their help in parental care and themselves becoming a resource (e.g. partner compensation;***Chen et al., 2020***; ***Eggert et al., 2008***; ***Eggert and Müller, 2000***; ***Royle and Hopwood, 2017***). The formation of such groups and associated social interactions are largely determined by constrained resources that limit each individual’s breeding opportunities (e.g. carcass availability; ***Eggert et al., 2008***; ***Eggert and Sakaluk, 2000***; ***Komdeur et al., 2013***; ***Müller et al., 2007***), and are also influenced by environmental conditions (***Chen et al., 2020***; ***Liu, Chan, et al., 2020***; ***Trumbo, 1992***). For example, in adverse environments, e.g. harsh and unpredictable climate and strong interspecific competition, within-group conflict among individuals is often low, and individuals are more likely to invest in cooperation (***Chen et al., 2020***; ***Liu, Chan, et al., 2020***). In such groups, winning individuals likely become dominants that monopolize the carcass after several rounds of fights, whereas subordinates have restricted access towards the carcass (***Eggert et al., 2008***; ***Eggert and Müller, 1992***; ***Müller et al., 2007***). Because reproductive conflict is more frequent among same-sex individuals, dominants always show different aggressive and parental behaviour towards the same- or opposite-sex subordinates (***Eggert and Müller, 1992***, ***2000***). In particular, dominant males often show more aggressive behaviour towards subordinate males than subordinate females (e.g. direct eviction from the carcass; ***Mitchell et al., 2009***). Due to the high costs for individuals of an escalated conflict (e.g. rejecting others from groups) and their inability to fully control the reproduction of others (***Eggert et al., 2008***; ***Hopwood et al., 2013***; ***Liu, Chan, et al., 2020***; ***Shen and Reeve, 2010***), this scenario may promote a tug-of-war competition with a tolerance interaction, in which dominants and subordinates reach a negotiated settlement (or a stable struggle equilibrium) in the allocation of resources and reproduction (***Cant and Young, 2013***; ***Clutton-Brock, 1998***; ***Eggert and Müller, 2011***; ***Gilchrist, 2006***). Such mutual tolerance is typically characterized by a low level of aggressiveness (e.g. increased access towards the carcass for subordinates) and no or moderate skewed reproduction between individuals (***Eggert et al., 2008***; ***Eggert and Müller, 2000***; ***Shen and Reeve, 2010***). Nevertheless, it remains unclear how such a negotiation or struggle affects an individual’s behaviour and reproduction, as well as the level of cooperation and even group productivity (e.g. reproductive success of groups; ***Cant, 2000***; ***Clutton-Brock, 1998***; ***Langer et al., 2004***; ***Reeve et al., 1998***). On the basis of the supposed tug-of-war competition model, this can be addressed from two perspectives (***Bradley et al., 2005***; ***Hamilton, 2013***; ***Langer et al., 2004***). From the subordinate’s point of view, it is important to study whether and how subordinates pay to stay in groups (or to reproduce concurrently with dominants). From the dominant’s point of view, it is crucial to study whether dominants pay costs from living with subordinates in groups, and if subordinates become a resource that compensates for these costs of group living or breeding. Addressing and integrating these perspectives will offer comprehensive insights into the evolution of social behaviour driven by a tug-of-war competition.

Here, we conducted experiments using the burying beetles, *N. vespilloides*, to test whether subordinates pay by helping in carcass preparation to enhance their fitness benefits and group productivity (‘paying-to-stay’), and whether dominants pay costs from living with subordinates (‘pay-from-staying’) in communal breeding (***Eggert and Müller, 2011***; ***Liu, Chan, et al., 2020***; ***Ratz and Smiseth, 2018***; ***Smiseth et al., 2005***; ***Fig. 1***). First, we manipulated the cooperative behaviour of subordinates of both sexes (i.e. subordinates helped or did not help dominants in carcass preparation), as well as the pressure of interspecific competition (i.e. the absence or presence of blowfly maggots that compete with beetles for carcasses for reproduction). We predict that subordinates that paid the costs by helping in carcass preparation may receive a high level of tolerance (e.g. increased parental investment time for individuals) and even direct reproductive benefits in groups (determined using parentage analysis), and that there occurs a reduced level of such tolerance towards subordinates under an observed high pressure of interspecific competition with blowfly maggots. Second, we conducted a removal experiment of dominants of either sex to investigate the behavioural responses of subordinates to the absence of dominants, and to examine whether remaining dominants pay more costs from living with subordinates in the group, namely sex difference in compensation behaviour to the loss of breeding partners. We expect that a single remaining dominant exhibits less aggressiveness towards same-sex subordinates and more parental investment (i.e. increased access towards the carcass and providing more care on the carcass) compared to opposite-sex subordinates in the absence of its dominant partner. Subordinates of the opposite sex may pay less costs in order to stay in the group due to the fact that they themselves become an important resource for the remaining dominant so as to increase fitness benefits (e.g. reproduce with the remaining dominant). We also expect that a single remaining dominant has a worse parenting performance and has less ability to control subordinates, and consequently may obtain reduced benefits (i.e. lower reproduction) from living with subordinates. Third, we examined the effects of the extent of a tug-of-war competition over parental investment between dominants and subordinates on group stability and overall group productivity. Given a trade-off between individual reproduction and group productivity in communal groups, we expect a negative correlation with respect to parental investment (e.g. parental investment time and weight change) between dominant and subordinate individuals. We also expect that such a correlation reflects the intensity of a within-group tug-of-war competition between individuals, which negatively influences group productivity (i.e. the higher the intensity of a within-group tug-of-war competition, the lower group productivity should be).

**Fig. 1.**
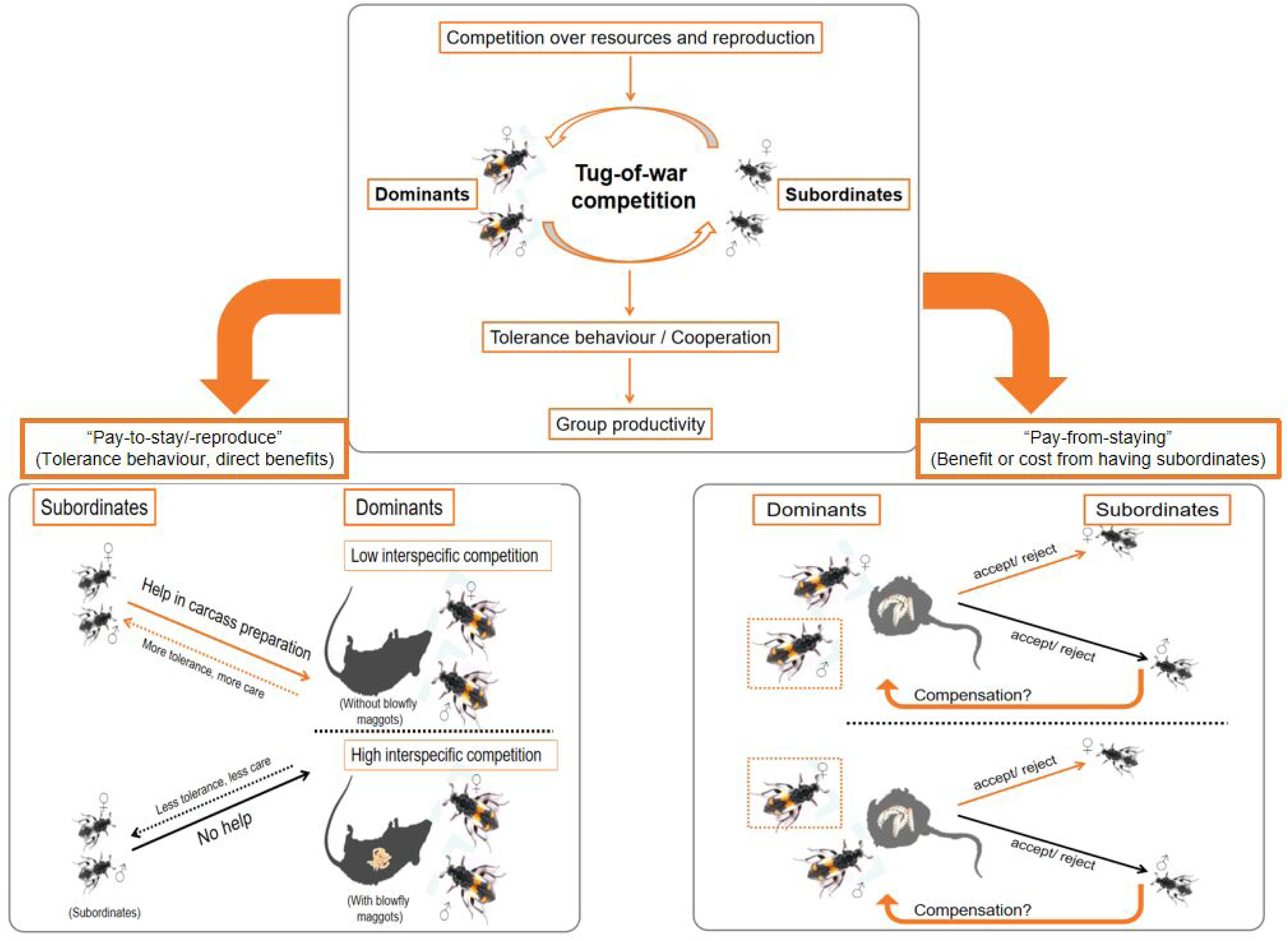
Schematic view of the experimental procedure in communal breeding of burying beetles, *N. vespilloides*. The experiment can be divided into two phases. In the manipulation experiment, the cooperative status of subordinates and the absence or presence of blowfly maggots were manipulated to test whether subordinates paid to stay on carcasses by helping dominants in carcass preparation (“pay-to-stay/-reproduce” hypothesis). In the removal experiment, either dominant females or males were removed, and the behavioural responses of the remaining dominant to subordinate females and males were tested (“pay-from-staying” hypothesis).

## Results

### Pay to stay: Effects of the cooperative status of subordinates and the pressure of interspecific competition with blowfly maggots on parental investment and reproductive success

To understand whether subordinates paid to stay by helping dominants in communal groups, and whether this was also influenced by interspecific competition, we conducted a manipulation experiment to examine the effects of the ‘cooperative behaviour’ status of subordinates (i.e. subordinates assisted dominants in carcass preparation or not) and the pressure of interspecific competition (i.e. absence or presence of blowfly maggots) on individual parental behaviour and investment and reproduction (***see supplementary materials: Fig. S1***). We found no effect of interspecific competition on individual dominant status in communally breeding groups, as large individuals were dominants that spent significantly more time on the carcass than small individuals (i.e. subordinates) in the absence or presence of blowfly maggots (z = 21.04, p < 0.001 and z = 5.36, p < 0.001; ***Fig. 2; see supplementary materials: Table S1***). However, the presence of blowfly maggots resulted in a significant reduction in the parental investment time for dominants (mean ± sd: 33.86 ± 2.92%, n = 33 vs. 67.75 ± 2.26%, n = 58; z = 5.02, p < 0.001) but not for subordinates (8.38 ± 1.72%, n = 33 vs. 8.16 ± 1.30%, n = 58; z = 1.66, p = 0.18) compared with the absence of blowfly maggots (***Table S1***). The ‘cooperative’ status of subordinates (i.e. non-cooperative, cooperative and pseudo-cooperative; whether or not subordinates assisted dominants with carcass preparation) determined the tolerance level of dominants towards subordinates (i.e. increased time spent providing care on the carcass for subordinates; ***Fig. 2a,b; Table S1***). In addition, such tolerance level was mediated by the pressure of interspecific competition with blowfly maggots (***Fig. 2a,b; Table S1***). Irrespective of the presence or absence of blowfly maggots, dominant individuals did not respond to the ‘cooperative’ status of subordinates by adjusting their amount of time providing care on the carcass (***Fig. 2a; Table S1***). In the absence of blowfly maggots, ‘cooperative’ subordinates that assisted dominants with carcass preparation had a significant increase in time spent on the carcass compared with ‘non-cooperative’ subordinates that did not help in carcass preparation (10.86 ± 2.28%, n = 24 vs. 4.04 ± 0.98%, n = 22; ***Fig. 2b; Table S1***), suggesting the occurrence of an increased tolerance level of dominants towards ‘cooperative’ subordinates compared with ‘non-cooperative’ subordinates. Such an increased level of tolerance in the absence of blowfly maggots was due to subordinates’ assistance in carcass preparation, as the amount of time spent on the carcass was similar between ‘cooperative’ and ‘pseudo-cooperative’ subordinates (i.e. subordinates did not jointly bury the carcass with dominants, but had buried another carcasses by themselves; 10.86 ± 2.28%, n = 24 vs. 10.32 ± 3.59%, n = 12; z = 0.56, p = 0.99), and between ‘non-’ and ‘pseudo-cooperative’ subordinates (4.04 ± 0.98% vs. 10.32 ± 3.59%; z = -2.73, p = 0.06; ***Fig. 2b; Table S1***). However, in the presence of blowfly maggots, the amount of time spent on the carcass was similar for subordinates that differed with respect to the ‘cooperative’ status (non-cooperative: 7.89 ± 3.32%, n = 10; cooperative: 11.54 ± 3.48%, n = 11; pseudo-cooperative: 6.15 ± 2.43%, n = 12; ***Fig. 2b; Table S1***). Additionally, in the presence of blowfly maggots, all individuals had a reduced time in total group investment compared with individuals in the control treatment without blowfly maggots (21.12 ± 1.44%, n = 33 vs. 39.28 ± 1.25%, n = 58; χ^2^ = 51.90, p < 0.001; ***Fig. 2c; Table S1***). However, in both the absence and presence of blowfly maggots, the total group investment time was similar among all groups for which subordinates differed with respect to their cooperative status (χ^2^ = 1.39, p = 0.50; ***Fig. 2c; Table S1***).

**Fig. 2.**
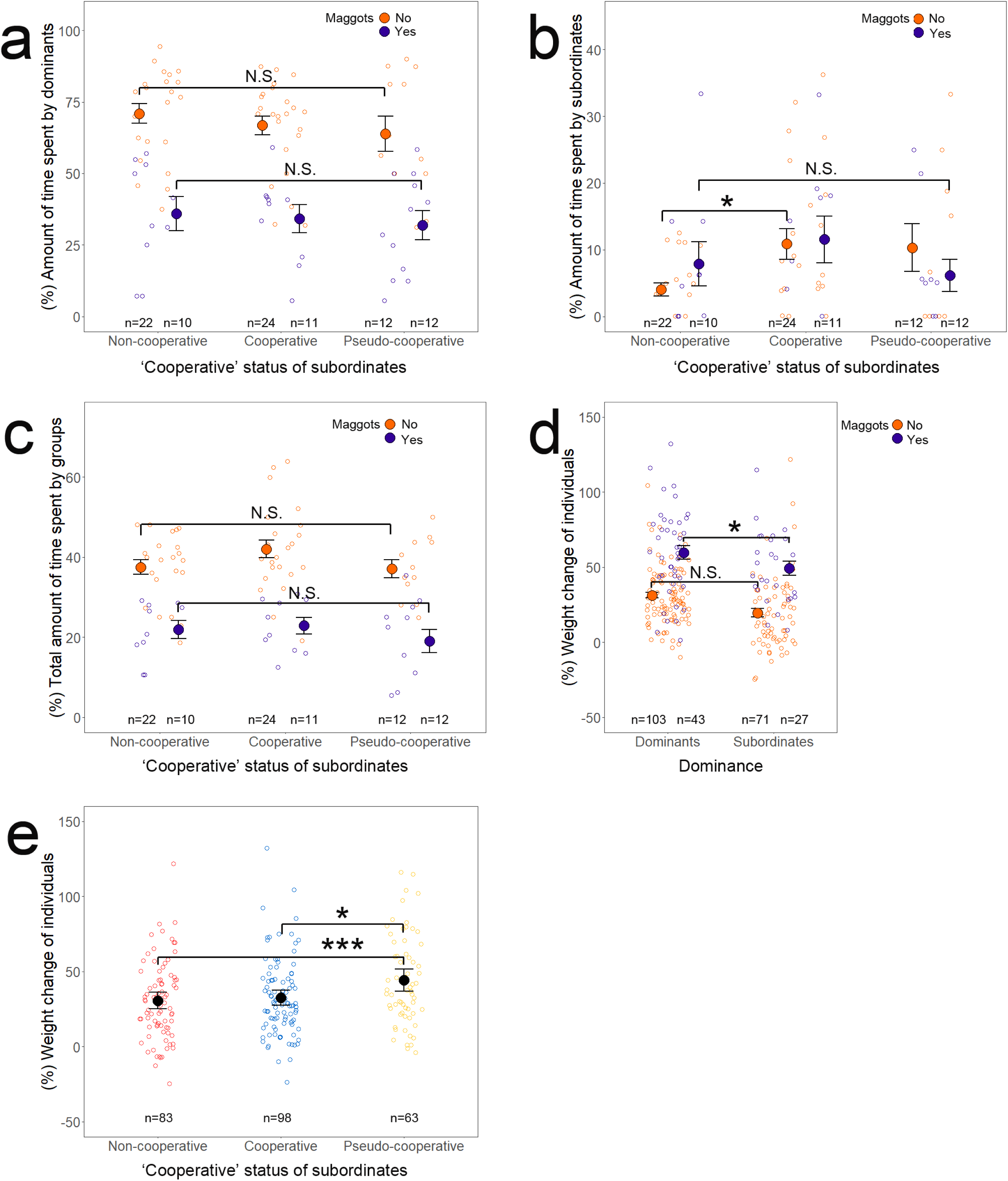
“Pay-to-stay”: Effects of the ‘cooperative’ status of subordinates and the pressure of interspecific competition with blowfly maggots on parental care in communal breeding of burying beetles, *N. vespilloides*. **(a-c)** Effects of the level of ‘cooperative’ status of subordinates on the amount of time spent by dominants (i.e. the dominant pair) **(a)** and subordinates (i.e. the subordinate pair) **(b)**, and on the total amount of time spent by groups **(c)** in the absence and the presence of blowfly maggots. **(d)** Effect of dominance status on weight change of individuals. **(e)** Effect of the level of ‘cooperative’ status of subordinates on weight changes of individuals. X-axis labels indicate the level of ‘cooperative’ status of subordinate individuals: non-cooperative, cooperative and pseudo-cooperative status, and the level of dominance status of individuals: dominants and subordinates. Blowfly maggots absence (maggots present: No; organe dots) and presence (maggots present: Yes; purple dots), and female (red dots) and male (blue dots) individuals. Notes: Asterisks above error bar indicate significance: *, p < 0.05; **, p < 0.01; ***, p < 0.001. Raw data and means ± standard errors (SE) are shown. See Table S1 for more statistical details.

In the absence of blowfly maggots, no difference in weight change was observed for both dominant and subordinate individuals (31.29 ± 1.91%, n = 103 vs. 19.64 ± 2.95%, n = 71; t = 2.24, p = 0.05), whereas dominants gained more weight than subordinates in the presence of blowfly maggots (59.74 ± 4.46%, n = 43 vs. 49.31 ± 4.65%, n = 27; t = 2.93, p = 0.01; ***Fig. 2d; Table S1***). Additionally, in the absence of blowfly maggots, dominant females gained more weight than dominant males (38.24 ± 2.79%, n = 54 vs. 23.66 ± 2.11%, n = 49; t = 2.81, p = 0.02), whereas no sex difference in weight change was observed for subordinates (20.93 ± 4.51%, n = 42 vs. 17.76 ± 3.14%, n = 29; t = -0.88, p = 0.85; ***see supplementary materials: Fig. S3; Table S1***). However, in the presence of blowfly maggots, no sex difference in weight change was observed for both dominants and subordinates (t = 1.93, p = 0.20 and t = 2.37, p = 0.07; ***Fig. S3; Table S1***). In groups with ‘pseudo-cooperative’ subordinates, both dominant and subordinate individuals gained more weight compared to individuals from other groups (non-cooperative: 30.66 ± 2.84%, n = 83; cooperative: 32.48 ± 2.59%, n = 98; pseudo-cooperative: 44.28 ± 3.79%, n = 63; t = 3.65, p < 0.001 and t = 2.62, p = 0.02; ***Fig. 2e; Table S1***).

The presence of blowfly maggots resulted in a significant reduction in the reproductive success of groups compared to when blowfly maggots were absent (with maggots, n = 18 vs. without maggots, n = 58; brood size: 14.00 ± 1.01 vs. 20.10 ± 1.33; mean larval weight: 0.1268 ± 0.0066g vs. 0.1486 ± 0.0047g; ***Fig. S4; Table S2***). The ‘cooperative’ status of subordinates differentially modulated the reproductive success in the absence or presence of blowfly maggots (***Fig. 3a,b; Table S2***). When subordinates were ‘cooperative’ in carcass preparation, groups produced lighter larvae in the presence of blowfly maggots than in the absence of maggots (0.1037 ± 0.0095g, n = 8 vs. 0.1462 ± 0.0073g, n = 24; z = 2.94, p = 0.01; ***Fig. 3b***). The brood size of groups was similar between the two treatments (14.12 ± 2.44 vs. 22.75 ± 2.39; z = 2.27, p = 0.08; ***Fig. 3a and S4; Table S2***). However, when subordinates were ‘non-’ or ‘pseudo-cooperative’, the absence or presence of blowfly maggots did not influence brood size and mean larval weight (***Table S2***). Although dominant individuals produced more offspring than subordinate individuals in groups with ‘cooperative’ or ‘non-cooperative’ subordinates, there were no significant differences in the number of offspring produced by ‘cooperative’ vs. ‘non-cooperative’ subordinate females (4.30 ± 1.470, n = 10 vs. 1.78 ± 1.16, n = 9) or males (1.10 ± 0.62, n = 10 vs. 1.56 ± 1.31, n = 9; ***Fig. 3c; Table S3***), as well as the percentage of offspring produced by cooperative vs. non-cooperative dominant pairs (79.13 ± 2.96%, n = 40 vs. 90.11 ± 2.64%, n = 36; ***Table S3***).

**Fig. 3.**
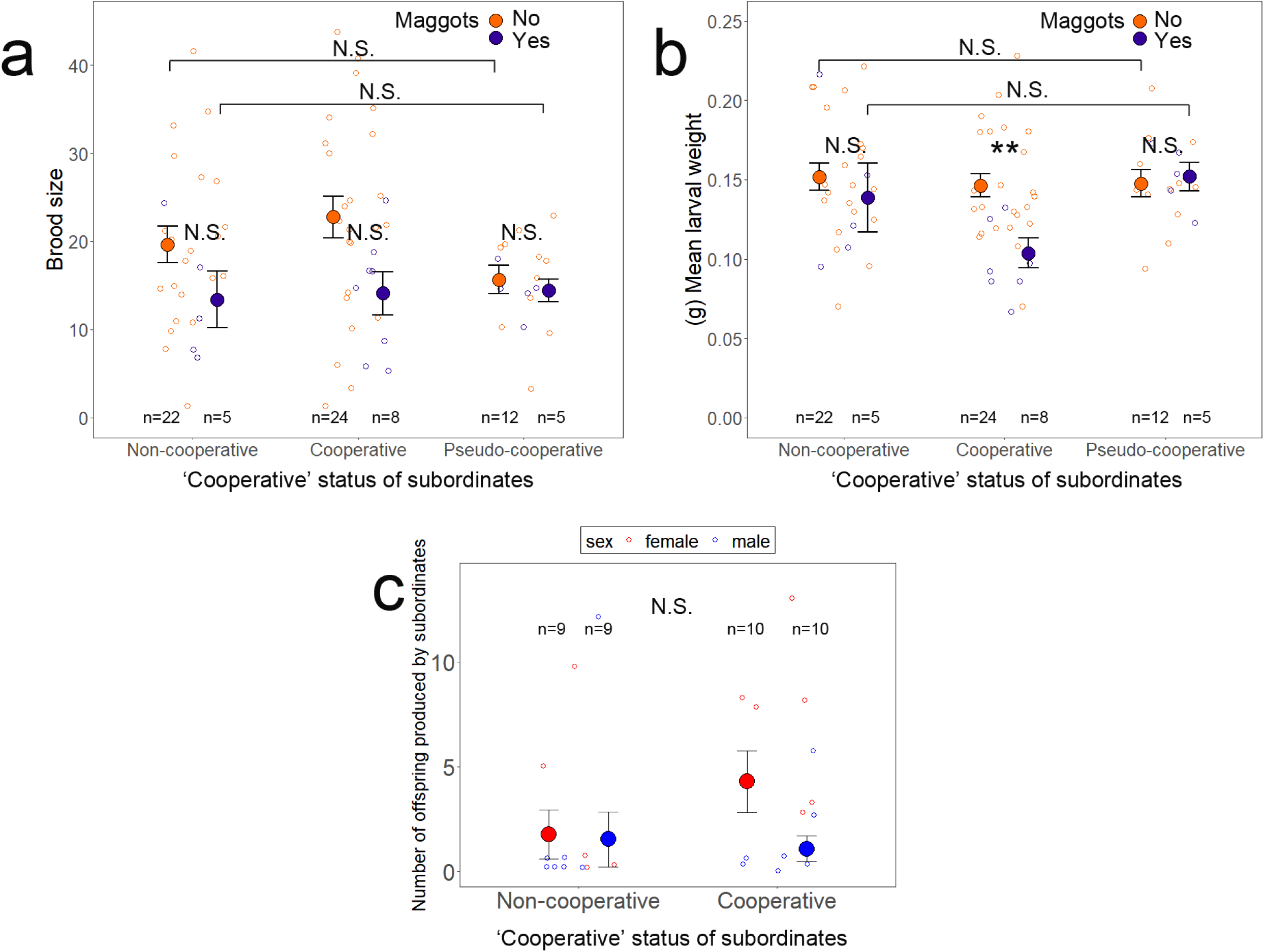
“Pay-to-stay”: Effects of the ‘cooperative’ status of subordinates and the pressure of interspecific competition with blowfly maggots on the reproductive success of groups and subordinate reproduction in communal breeding of burying beetles, *N. vespilloides*. **(a,b)** Effects of the level of ‘cooperative’ status of subordinates on brood size **(a)** and mean larval weight **(b)** of groups in the absence and presence of blowfly maggots. **(c)** Effect of the level of ‘cooperative’ status of subordinates on the number of offspring produced by female and male subordinates. X-axis labels indicate the level of ‘cooperative’ status of subordinate individuals: non-cooperative, cooperative and pseudo-cooperative statuses. Blowfly maggots absence (maggots present: No; orange dots) and presence (maggots present: Yes; purple dots), and female (red dots) and male (blue dots) individuals. Notes: Asterisks above error bar indicate significance: *, p < 0.05; **, p < 0.01; ***, p < 0.001. Raw data and means ± standard errors (SE) are shown. See Table S2 and S3 for more details.

### Subordinates respond to the removal of a dominant

To study whether subordinates had differential responses in parental behaviour and investment towards the absence of a dominant, we experimentally removed either a dominant female or male from communal groups, and tested the effects of these removals on individual parental behaviour and investment and reproduction (***Fig. 4 and 5; Table S4 and S6***). We also removed both dominant individuals from communal groups to verify whether the presence of a carcass, but not direct exclusion by dominants, influenced individual subsequent parental behaviour. Our results showed a sex difference in the behavioural response of dominants towards subordinates of the same versus opposite sex. Specifically, in communally breeding groups, the removal of either a dominant female or male had no significant influence on the individual time spent providing care on the carcass by subordinate females and males (***Fig. 4a,c; Table S4***). Subordinate females responded to the removal of both dominants by significantly increasing their amount of time spent on the carcass compared with the removal of a dominant male or female and without the removal of dominants (removal of both dominants: 39.05 ± 8.24%, n = 16; of a dominant male: 7.56 ± 3.67%, n = 19; of a dominant female: 17.24 ± 5.43%, n = 19; without removal of dominants: 14.39 ± 3.75%, n = 20; z = 3.80, p = 0.002 and z = 3.09, p = 0.02; ***Table S4***), whereas this was not the case for subordinate males (***Fig. 4a,c; Table S4***). At the pair level, subordinate individuals had similar parental investment time on the carcass between groups with either the dominant females or males remove, or without the removal of dominants. However, a significantly higher parental investment by subordinate pairs was observed when both dominants had been removed compared with when no dominants were removed (12.50 ± 2.80%, n = 20 vs. 31.25 ± 5.47%, n = 16; ***Fig. 5a; Table S4***). Subordinates did not adjust their parental investment in response to the absence of dominant females or males, but they increased their parental investment time when both dominants were absent (***Fig. 5c; Table S4***). Our results also showed that the removal of a dominant female resulted in a greater cost in parental investment compared to the removal of a dominant male (***Fig. 5c; Table S4***). The removal of dominant females, but not the removal of dominant males, had a significant reduction in the total group investment time compared with groups without the removal of dominants (removals of dominant females: 18.15 ± 1.77%, n = 19; of dominant males: 25.66 ± 2.31%, n = 19; without removal: 31.28 ± 2.70%, n = 20; ***Fig. 5c; Table S4***). Also, the removal of dominant females resulted in a significant reduction in the total group investment time compared with the removal of dominant males (Fig. 5c; Table S4). Groups with both dominants removed had a significantly lower total amount of time spent on the carcass compared with groups with dominant males removed and without the removal of dominants (both removal: 15.62 ± 2.74%, n = 16; ***Table S4***), suggesting that an increase in parental investment by subordinates could not fully compensate for the costs due to the loss of both dominants. Subordinate’s weight change was similar for both males and females (F_1,53_ = 3.31, p = 0.07) and was not influenced by the removal experiments (***Table S4***), while subordinate individuals gained fewer or lost more weight with larger brood sizes (F_1,53_ = 5.74, p = 0.02; ***Table S4***). Our results demonstrated that the loss of dominant females and males had no significant influence on group productivity, which may be due to subordinates compensating for the reproductive costs, as brood size of offspring and mean larval weight at larval dispersal were similar for the removal experiments (***Fig. 6a,b; Table S6***).

**Fig. 4.**
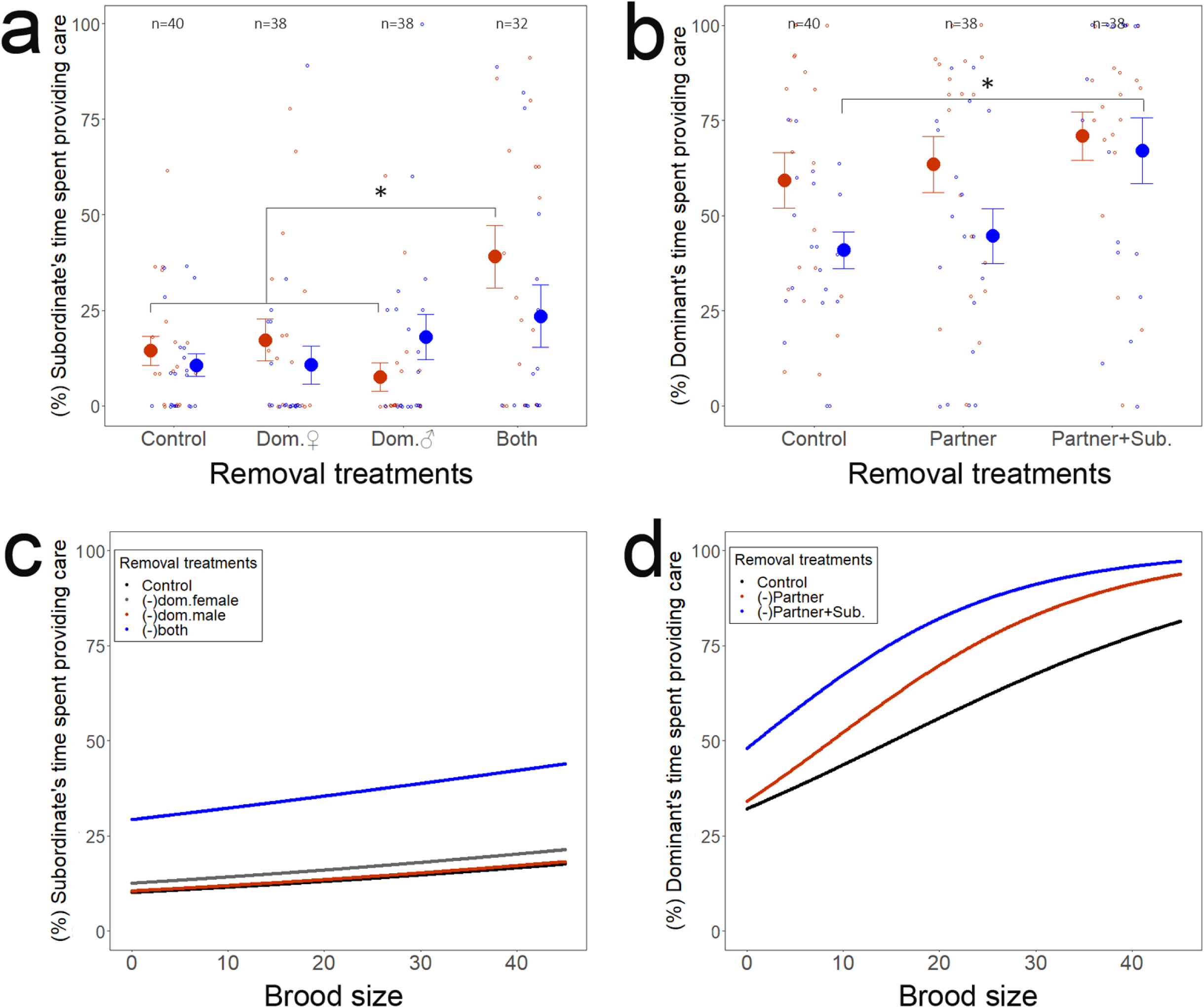
Effects of removal treatments on subordinates and dominants in parental care. **(a and c)** Effects of the removals of dominant females and males **(a)**, and brood size of offspring **(c)** on the amount of care provided by subordinate females and males. X-axis labels indicate the removal of dominants: control (without removal of dominant individuals; black line), dom.♀ (removal of dominant female; grey line), dom.♂(removal of dominant male; red line), and both (removal of both dominant individuals; black line). **(b and d)** Effect of the removal of dominant partners and subordinates **(b)**, and brood size of offspring **(d)** on the amount of care provided by dominant females and males. X-axis labels indicate the removals of dominant partners and subordinates: control (no removals of dominants and subordinates; black line), partner (removals of dominant female or male; red line), and partner+sub. (removals of one of dominants and subordinates; blue line). Notes: Asterisks above error bar indicate significance: *, p < 0.05; **, p < 0.01; ***, p < 0.001. Raw data and means ± standard errors (SE) are shown. In **(a and b)**, dominant and subordinate female (red dots) and male individuals (blue dots). Numbers in the graph indicate sample sizes: n = number of individuals. See Table S4 and S5 for more statistical details.

**Fig. 5.**
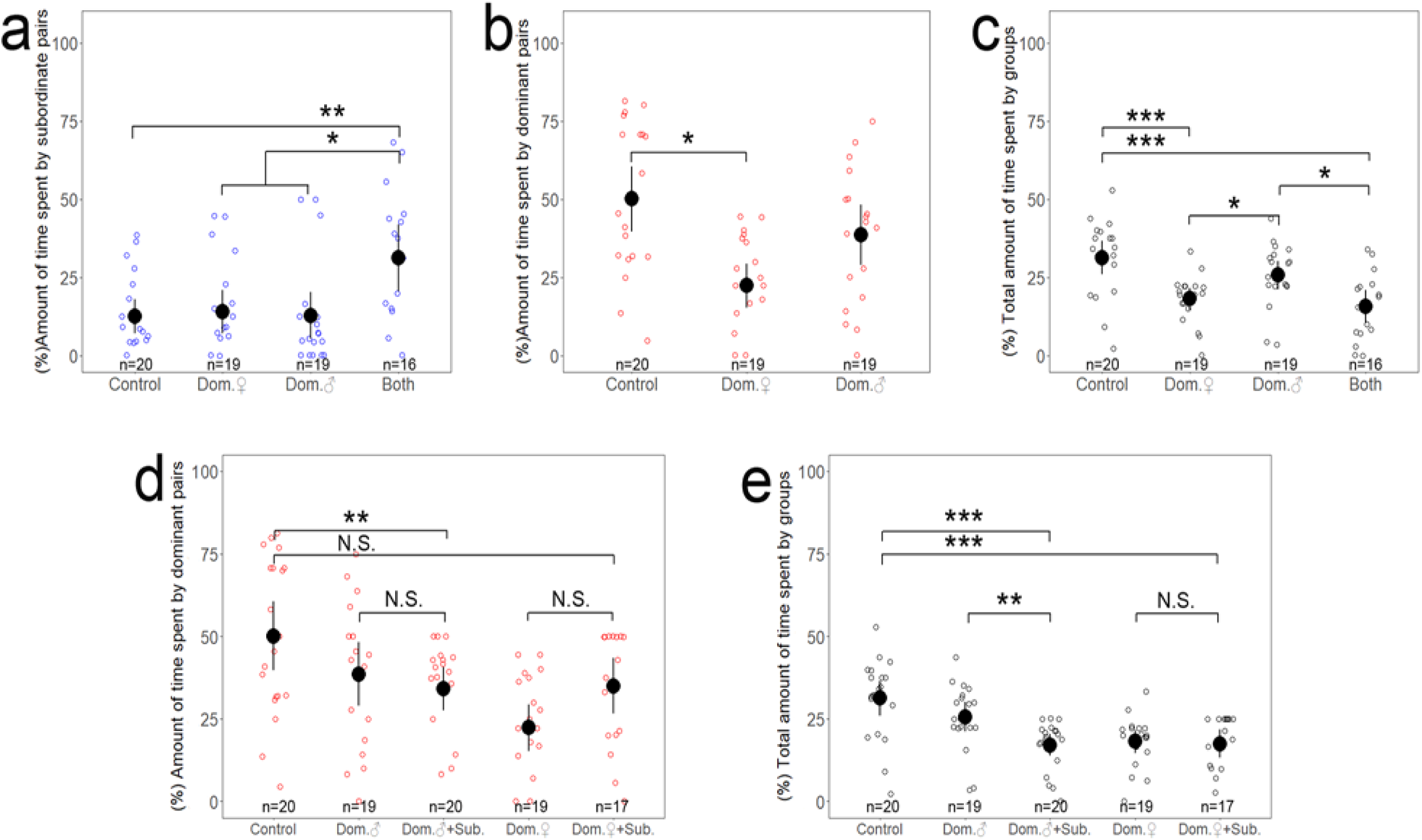
Effects of removal treatments on subordinates and dominants in parental care. **(a and c)** Effects of the removals of dominant females and males on amount of time spent providing care by subordinates **(a)** and by dominants **(b)**, and on the total amount of time spent by groups **(c). (d and e)** Effect of the removal of dominant partners and subordinates on the amount of time spent providing care by dominants **(d)**, and the total amount of time spent by groups **(e)**. X-axis labels indicate that the removals of dominant partners and subordinates: control (without removal of dominants and subordinates), dom.♂ (removal of dominant male) and dom.♂+sub. (removal of dominant male and subordinates), dom.♀(removal of dominant female) and dom.♀+sub. (removal of dominant female and subordinates) and both (removal of both dominant individuals). Notes: Asterisks above error bar indicate significance: *, p < 0.05; **, p < 0.01; ***, p < 0.001. Raw data and means ± standard errors (SE) are shown. See Table S6 for more details.

**Fig. 6.**
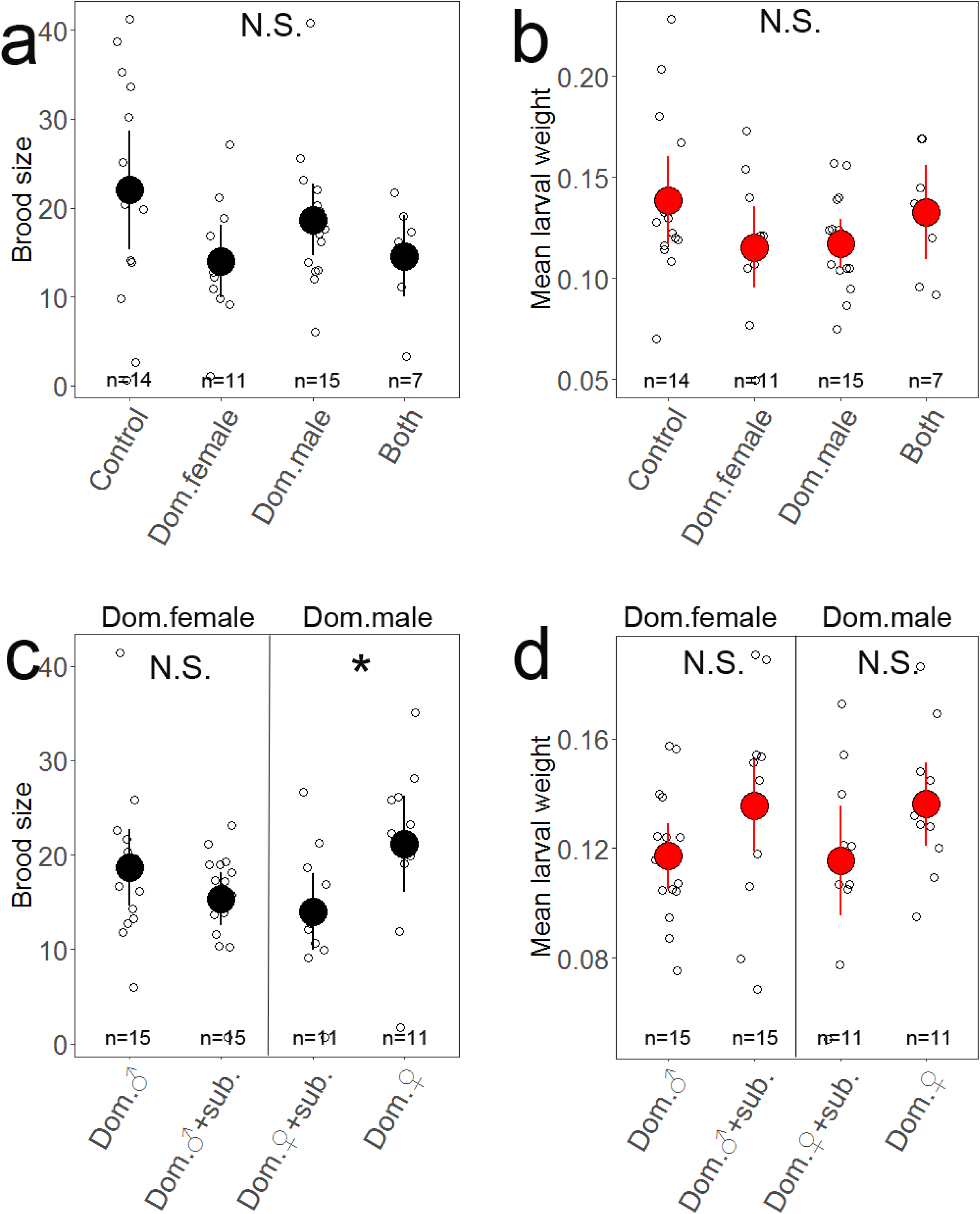
Effects of removal treatments of dominants and subordinates on the reproductive success of groups. **(a and b)** Effects of the removals of dominant females and males on **(a)** brood size, and **(b)** mean larval weight of groups at larvae-dispersal. X-axis labels indicate the removals of dominants: control (without removal of dominants), dom. female (removal of dominant female), dom.male (removal of dominant male), and both (removal of both dominant individuals). **(c and d)** Effects of the removals of dominant partners and subordinates on **(c)** brood size, and **(d)** mean larval weight of groups at larvae-dispersal. X-axis labels indicate the removals of dominant partners and subordinates: dom.♂ (removals of dominant male) and dom.♂+sub. (removals of dominant male and subordinates) for dominant females; dom.♀ (removals of dominant females) and dom.♀+sub. (removals of dominant females and subordinates) for dominant males. Notes: Asterisks above error bar indicate significance: *, p < 0.05; **, p < 0.01; ***, p < 0.001. Raw data and means ± standard errors (SE) are shown. See Table S7 and S8 for more details.

### Pay-from-staying: Remaining dominants respond to the removals of their partners and subordinates

To examine whether a single remaining dominants compensated for the absence of its dominant partner in the absence of subordinates and paid more costs from the presence of subordinates in communal groups, we removed either a dominant female or male, as well as both subordinates, and tested the effects of these removals on individual parental behaviour and investment and reproduction (***Fig. 4 and 5; Table S5***). Our results showed experimental evidence for a sex difference in compensation behaviour for the loss of partners by dominant individuals, and such compensation in parental investment was also affected by the presence of subordinates (***Fig. 4b,d and 5d,e; Table S5***). The remaining dominant females did not respond to the removal of their dominant partners by not adjusting their parental investment time in the presence or absence of subordinates (***Fig. 4b,d; Table S5***). Compared with groups without the removal of dominants, the removal of dominant males did not influence the investment time by dominant pairs in the presence of subordinates (38.56 ± 4.91%, n = 19 vs. 50.07 ± 5.32%, n = 20), but resulted in a significant reduction in the amount of time spent by dominant pairs when subordinates were absent (17.06 ± 1.65%, n = 20 vs. 50.07 ± 5.32%, n = 20; ***Fig. 5d; Table S5***). These findings suggested that the remaining dominant females fully compensated for a reduced parental investment due to the loss of their partners in the presence of subordinates, but they showed partial or no compensation in parental investment when subordinates were absent. However, the parental investment level by dominant males was significantly lower in the presence of subordinates than in the absence of subordinates (38.56 ± 4.91%, n = 19 vs. 50.07 ± 5.32%, n = 20; ***Fig. 4b,d and 5d; Table S5***). Furthermore, our results showed that the presence of dominant females could compensate for the loss of their dominant partners in parental investment, as groups without the removal dominants or with dominant males removed had similar total group investments in parental care (31.28 ± 2.70%, n = 20 vs. 25.66 ± 2.31%, n = 19; ***Fig. 5d; Table S5***). Compared with these two groups, there was a significant reduction in the total parental investment of groups when dominant males and subordinates were both removed (17.06 ± 1.65%, n = 20; ***Fig. 5e; Table S5***), which indicated an important role of subordinates in contributing to the total parental investment of groups.

The remaining dominant males did not compensate for a reduced parental investment due to the loss of their dominant partners in the presence of subordinates, but they compensated for such loss by increasing their investment when subordinates were absent (***Fig. 4b,d and 5d,e; Table S5***). Specifically, compared with groups without the removal of dominants (40.91 ± 4.86%, n = 20), the remaining dominant males did not respond to the removal of their dominant partners by adjusting their amount of time spent on the carcass in the presence of subordinates (44.68 ± 7.20%, n = 19), but they significantly increased their amount of time on the carcass in the absence of subordinates (70.00 ± 8.60%, n = 17; ***Fig. 4b,d; Table S5***). Compared with no removal of dominants (50.07 ± 5.32%, n = 20), the removal of dominant females resulted in a significant reduction in the amount of time spent by dominant pairs in the presence of subordinates (22.34 ± 3.60%, n = 19) but had no effect on the amount of time spent by dominant pairs when subordinates were absent (17.50 ± 2.15%, n = 17; ***Fig. 5d; Table S5***). These results highlighted the key role of dominant females because they contributed more to parental investment than dominant males. The remaining dominant males did not fully compensate for the loss of their partners by increasing their level of parental investment or from living with subordinates in the group. Groups with the dominant female removed had a lower total group investment in parental care in the presence and absence of subordinates (19.15 ± 1.77%, n = 19 and 17.50 ± 2.15%, n = 17) compared with the control groups without the removal of dominants (31.28 ± 2.70%, n = 20; ***Fig. 5e; Table S5***).

Both the dominant females and males increased their amount of time spent on the carcass as brood size produced by groups increased (F_1,114_ = 35.64, p < 0.001; ***Fig. 4d; Table S5***). No significant differences in weight change of dominants were found when their partners and/or subordinates were removed (F_2,62_ = 1.24, df = 2, p = 0.30; ***Table S5***). The dominant males gained more weight than the dominant females when their dominant partners and subordinates were both removed (114.70 ± 73.42%, n = 16 vs. 37.52 ± 4.14%, n = 19; t = -2.81, p = 0.02; ***Table S5***). Groups with dominant females removed produced less offspring at larval dispersal when subordinates were present compared to when subordinates were absent (14.00 ± 2.08, n = 11 vs. 21.18 ± 2.61, n = 11; χ^2^ = 4.64, p = 0.04; ***Fig. 6a,c; Table S7***), while mean larval weight was similar (0.1154 ± 0.0102g vs. 0.1362 ± 0.0078g; F = 2.60, p = 0.12; ***Fig. 6d; Table S7***). When dominant males were removed, the reproductive success of groups was similar in the presence and absence of subordinates (***Fig. 6c,d; Table S7***).

### Tug-of-war competition between dominants and subordinates and its impact on group productivity

In communal groups of burying beetles, we observed a negative correlation for the amount of time spent on the carcass between dominant and subordinate pairs. Subordinate individuals reduced their time providing care and had fewer gains in weight (mean ± sd: 28.54 ± 2.57%, n = 114), if dominant individuals spent more time providing care (***Fig. 7a,c; Table S8 and S10***). However, the highest total amount of care provided by communal groups was found when both dominants and subordinates increased their time providing care on the carcass (***Table S8 and S10***). Independent of the subordinates’ cooperative status, all individual group members had fewer gains in weight (mean ± sd: 35.00 ± 1.31%, n = 277), if there was a higher total group investment in care (***Fig. 7b; Table S8 and S10***). In the absence or presence of blowfly maggots, dominant individuals gained less weight or lost more weight with an increase in the amount of time that pairs spent on the carcass (mean ± sd: 39.51 ± 1.98%, n = 163), while subordinate individuals significantly gained more weight by spending more time on the carcass (***Fig. 7c,d; Table S8 and S10***). In the absence of blowfly maggots, dominant individuals gained less weight (mean ± sd: 32.26 ± 1.74%, n = 120) with an increase in the amount of time spent on the carcass, while subordinates significantly gained more weight by spending more time on the carcass (mean ± sd: 59.74 ± 4.46%, n = 43; ***Fig. 7e,f; Table S8 and S10***). However, this was not the case for dominant and subordinate individuals in the presence of blowfly maggots (***Fig. 7e,f; Table S8 and S10***). At larval dispersal, an enhanced group productivity was found when dominant individuals spent more time providing care, which was the case in both the absence and presence of blowfly maggots (***Fig. 8a-c; Table S9 and S10***). Groups with dominants and subordinates that gained less weight produced larger brood sizes, while mean larval weight was not associated with their weight change (***Fig. 8d-f; Table S9 and S10***).

**Fig. 7.**
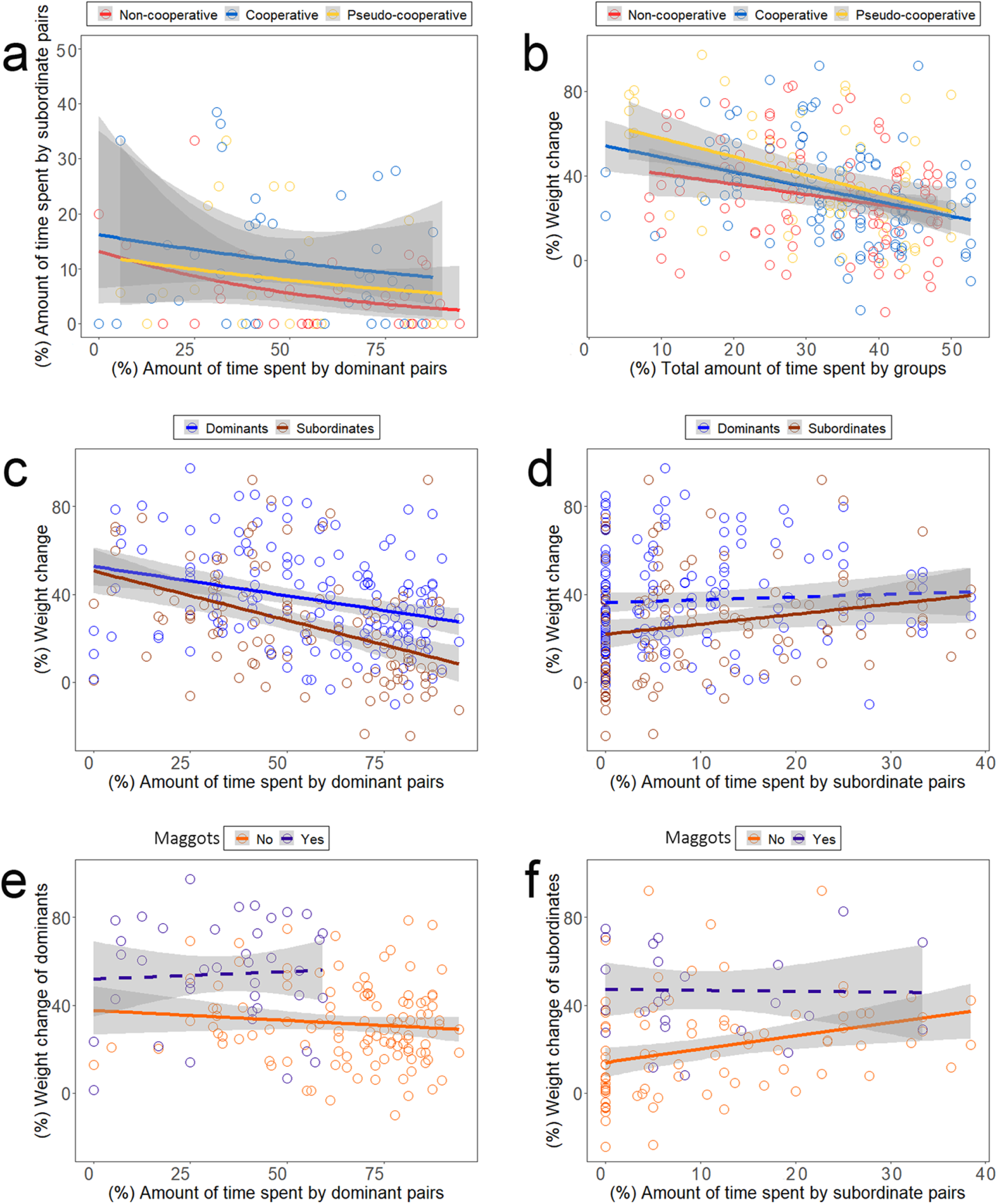
“Tug-of-war” competition between dominants and subordinates in communally breeding burying beetles, *N. vespilloides*. **(a)** Relationship of the amount of time spent providing care on the carcass by subordinate pairs with the amount of time spent by dominant pairs. **(b)** Relationship of weight change of individuals with the total amount of time spent by groups (dominant and subordinate pairs combined). **(c and d)** Relationships of weight change of dominant and subordinate individuals with the amount of time spent by **(c)** dominant pairs, and by **(d)** subordinate pairs. **(e and f)** Relationships of weight change of dominant individuals with the amount of time spent by dominant pairs **(e)**, and of weight change of subordinate individuals with the amount of time spent by subordinate pairs **(f)**, in the absence and the presence of blowfly maggots. Notes: The level of ‘cooperative’ status of subordinate individuals: non-cooperative (red lines and dots), cooperative (blue lines and dots) and pseudo-cooperative statuses (yellow lines and dots). Dominant (red lines and dots) and subordinate individuals (blue lines and dots) with significant (solid lines) or non-significant (dashed lines) correlations. Blowfly maggots absence (orange lines and dots) and presence (purple lines and dots) with significant (solid lines) or non-significant (dashed lines) correlations. See Table S9 for more statistical details.

**Fig. 8.**
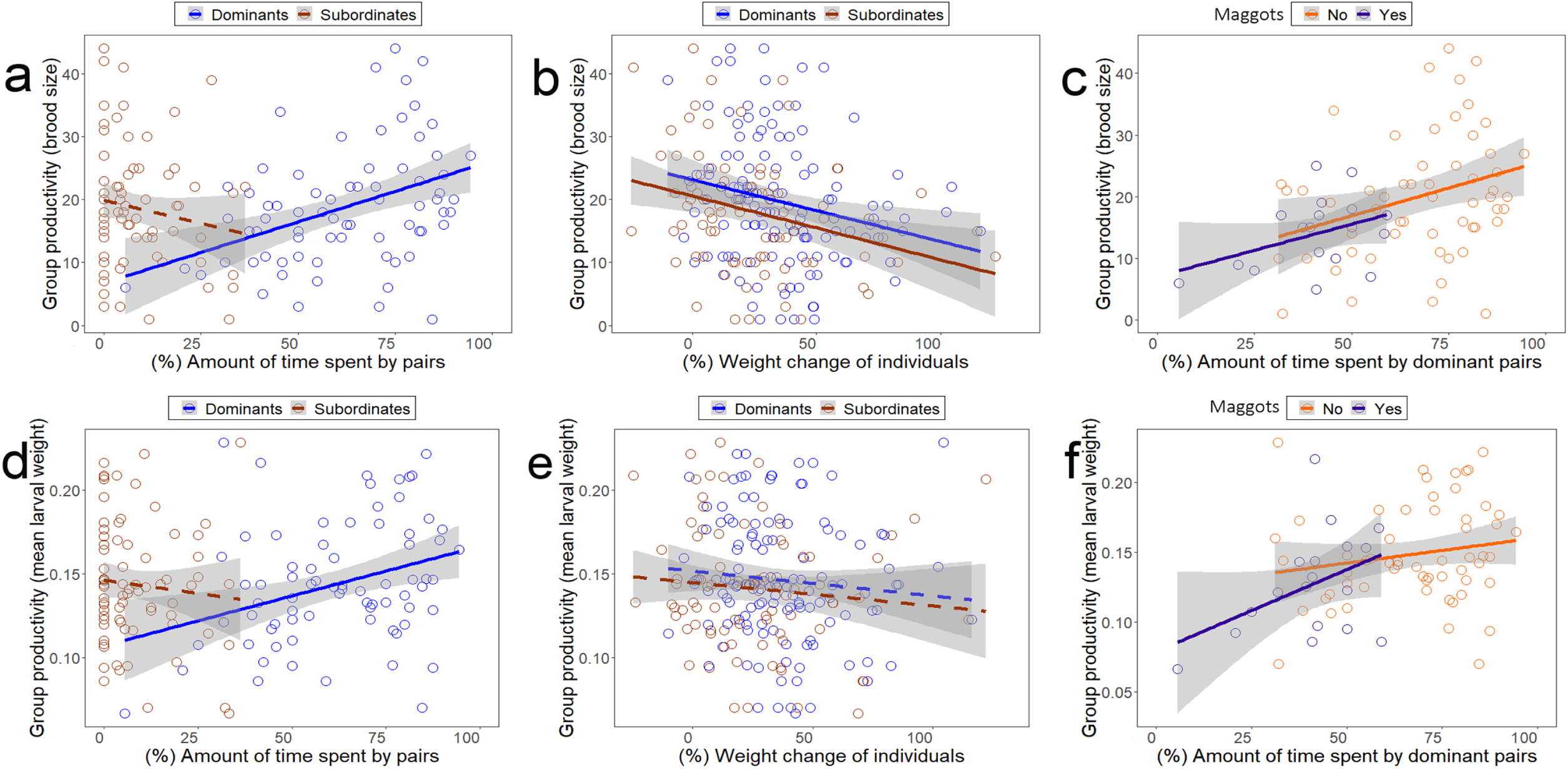
Effects of the amount of parental care provided by dominant and subordinate pairs and individual weight on group productivity (i.e. the reproductive success of groups) in communally breeding burying beetles, *N. vespilloides*. Relationships of the amount of time spent by dominant and subordinate pairs **(a and d)**, of weight changes of dominant and subordinate individuals **(b and e)**, and of the amount of time spent by dominant pairs in the absence and presence of blowfly maggots **(c and f)** on brood size **(a-c)** and mean larval weight of groups **(d-f)**. Notes: Dominant (red lines and dots) and subordinate individuals (blue lines and dots) with significant (solid lines) or non-significant (dashed lines) correlations. Blowfly maggots absence (orange lines and dots) and presence (purple lines and dots) with significant (solid lines) or non-significant (dashed lines) correlations. See Table S10 and S11 for more statistical details.

## Discussion

### Subordinates pay to stay in groups within interspecific competition

Our results show experimental evidence for the pay-to-stay hypothesis in communally breeding burying beetles, where subordinates that assist dominants with carcass preparation have an increased access towards the carcass (i.e. pay to be tolerated within a group), compared to subordinates that do not help in carcass preparation (***Fig. 2; supplementary materials: Table S1***). Such tolerance among group members represents the observed level of within-group conflict and is modulated by the pressure of interspecific competition (***Fig. 2; Table S1***). Specifically, there occurs a reduced tolerance of dominants towards subordinates in the high pressure of interspecific competition (with blowfly maggots) given the subordinate’s cooperative behaviour. This is probably due to an increased within-group tug-of-war competition between dominants and subordinates (***Gilchrist et al., 2004***; ***Langer et al., 2004***; ***Liu, Chan, et al., 2020***; ***Shen and Reeve, 2010***; ***Young et al., 2006***). Helping in parental care is an altruistic behaviour, for example, synergetic efforts in carcass preparation and alloparental care in burying beetles, as it imposes various costs on the donor, such as energy expenditure and sacrifices of other reproductive opportunities (***Billman et al., 2014***; Creighton et al., 2009; ***Duarte et al., 2018***; ***Scott, 1998***). We suggest that such cooperative services in parental investment may have evolved to enhance the extent of tolerance among potential communal breeders as they both can benefit from a low within-group conflict (***Eggert et al., 2008***; ***Heg and Hamilton, 2008***; ***Kokko et al., 2002***; ***Taborsky et al., 2016***). In many animal groups, ‘lazy-working’ or ‘free-riding’ individuals (i.e. they do not work as hard as other individuals in the group) are often tolerated by high-ranked individuals because they may become potential resources so as to enhance overall group benefits (***Breed et al., 1990***; ***Guindre-Parker and Rubenstein, 2018***; ***Hasegawa et al., 2016***; ***Kokko et al., 2002***; ***Liu, Chan, et al., 2020***). For example, under unfavourable environmental conditions, each individual can obtain enhanced synergetic group benefits from the presence of other members, and there occurs a low within-group conflict. In this scenario, each group members increase their investment in cooperation or decrease efforts in conflict because the cost of intraspecific competition increases as resource availability becomes limited or interspecific competition increases (***Liu, Chen, et al., 2020***; ***Shen et al., 2012***). However, the marginal costs of cooperation are different for dominant and subordinate individuals (***Johnstone and Cant, 1999***; ***Liu, Chen, et al., 2020***). In our study, dominant individuals are found to reduce their investment time in parental care and gain more weight when high blowfly competition is present, whereas all beetles that breed in communal groups fail to receive enhanced synergetic benefits of groups and more reproductive success (***Fig. 2 and 3; Table S2***). This could be explained by the marginal costs of cooperation exceeding the benefits of increasing total cooperative investment in groups when blowfly competition is very high. Also, it is likely that burying beetles rapidly adjust their strategies to decrease their own reproductive costs, based on the trade-off between investment in cooperation and individual benefits, because they may feed on the maggots as a preferred diet for energetic saving rather than invest more in shared efforts (***Liu, Chen, et al., 2020***; ***Müller et al., 2007***; ***Trumbo and Robinson, 2004***). For burying beetles, such a tolerance interaction seems not to require complex recognition, but may rely on direct benefits in resource use and reproduction (***Heg et al., 2009***; ***Jennions and Macdonald, 1994***; ***Kokko et al., 2002***; ***Ridley et al., 2004***; ***Steiger et al., 2008***). Helpful subordinates that invest in shared services (i.e. joint efforts in carcass preparation and defence) and own reproduction both receive high tolerance from others, which may generate generalized benefits of helping (***supplementary materials; Fig. S5; Table S11 and S12***). Different from direct reciprocity, such a generalized benefit is thought to not require individual recognition, or a scoring system or ‘biological markets’ to assess reputation(***Bshary and Noe, 2003***; ***Johnstone and Bshary, 2002***; ***Ridley et al., 2004***; ***Stevens and Gilby, 2004***). Therefore, we suggest that individuals may contribute to some common good, such as building a communal nest in burying beetles or other species, from which all group members subsequently obtain generalized benefits in reproduction (***Hamilton and Taborsky, 2005***; ***Heg et al., 2009***; ***Steiger et al., 2008***; ***Stevens and Gilby, 2004***).

We found no evidence that subordinates pay by helping in exchange for more reproductive benefits in shared broods, i.e. paying-to-reproduce (***Fig. 3; Table S3***). Although large individuals (i.e. dominants) frequently produce a large proportion of offspring in communal groups, the extent to which reproduction is partitioned between individuals is mainly dependent on carcass size (***Eggert et al., 2008***; ***Eggert and Müller, 1992***; ***Richardson and Smiseth, 2020***). Each cobreeding female is able to lay some eggs surrounding the carcass (***Eggert and Müller, 2000***, ***2011***; ***Müller et al., 1990***, ***2007***; ***Scott, 1998***), whereas each individual lacks the ability to recognize its own offspring and manipulate its reproductive shares, which may reduce the probability of committing infanticide, especially after larvae hatch (***Eggert et al., 2008***; ***Eggert and Müller, 2000***, ***2011***; ***Richardson and Smiseth, 2020***). Thus, it is likely that such reproductive partitioning is skewed before larvae-hatching, and during the egg-laying period. Subordinates may gain immediate and long-term benefits that are paid for by providing care, while such benefits are not easily distinguished in many group-living species (***Armitage and Schwartz, 2000***; ***Heg et al., 2009***). For example, in cooperatively breeding cichlids, subordinate females pay with alloparental care to get access to the breeding shelter, which may allow them to increase their opportunities to gain parentage (***Heg et al., 2009***; ***Werner et al., 2003***). Our results are not completely consistent with these findings and suggest that subordinates that pay with help in carcass preparation get access to the carcass or offspring, by which they may gain other direct fitness benefits, such as feeding from the carcass and sexual attractiveness (***Chemnitz et al., 2017***; ***Hopwood et al., 2013***; ***Richardson and Smiseth, 2020***; ***Wang et al., 2021***). Furthermore, it can be expected that access to resources for fitness benefits is exchanged reciprocally through mutualistic benefits between group members, in which subordinates pay with help in cooperative defence of resources against competitors, as well as communal nesting (***Clutton-Brock, 1998***; ***Liu, Chen, et al., 2020***; ***Zöttl, Heg, et al., 2013***). The ‘pay-to-stay’ strategy may occur to solve the optimal allocation of parental and alloparental care in groups, as well as the trade-off in efforts between cooperation and conflict (***Heg et al., 2009***; ***Johnstone and Cant, 1999***; ***Liu, Chen, et al., 2020***).

### Sex differences in the compensatory strategy for the loss of group members

Our results demonstrate a sex difference in subordinate’s parental behaviour in the absence of one of the dominants (***Fig. 4; Table S4***), which is due to dominants behaving differently towards same-versus opposite-sex subordinates (***Eggert and Müller, 2000***; ***Ratz et al., 2020***). We found that dominant females were more likely to reject subordinate females from the carcass if their dominant mates were removed compared with dominant males. However, subordinate males have similar parental investment when either dominant females or males were removed. Since subordinates staying in groups generates different costs for dominants due to intense within-group conflict (***Heg and Hamilton, 2008***; ***Mitchell et al., 2009***; ***Müller and Manser, 2007***; ***Trumbo, 2007***), we suggest that dominant females and males differ in their individual resource defence effort, which depends on the sex of the subordinates. Dominant individuals always show more aggressive and less affiliative behaviour towards same-sex subordinates, but not towards opposite-sex subordinates, because reproductive conflict is more frequent among same-sex individuals (***Eggert and Müller, 1992***, ***2000***; ***Mitchell et al., 2009***; ***Müller and Manser, 2007***). According to sex differences in costs and benefits of parental care and mating strategy (e.g. promiscuity), dominant females could obtain direct benefits in reproduction by successfully defending the resources, whereas males that have more mating opportunities may benefit from the presence of other females (***Eggert, 1992***; ***Ford and Smiseth, 2016***; ***Müller et al., 2007***; ***Smiseth and Moore, 2004***; ***Walling et al., 2009***). Therefore, we suggest that dominant females, but not dominant males, are expected to invest more in carcass defence and therefore restrict subordinate’s access towards the carcass (***Heg and Hamilton, 2008***; ***Mitchell et al., 2009***). Also, the amount of care provided by subordinate females is associated with their intrinsic motivations to stay within groups, because caring individuals could gain direct (e.g. higher maternity) or indirect benefits (e.g. breeding and contest experiences, saving resources for the future) by actively increasing access to the carcass (***Eggert et al., 2008***; ***Eggert and Müller, 1992***; Shippi et al., 2018; ***Smiseth et al., 2005***).

Our results show that dominant individuals deploy sex differences in the compensation strategy for the loss of mates (***Fig. 4 and 5; Table S4 and S5***). Specifically, the remaining female dominants may have no adjustment in parental investment in response to the loss of their mates, whereas the remaining male dominants only increase their parental investment when both their mates and subordinates are absent. Such sex difference regarding parental care can be explained by the fact that dominant females and males may deploy different strategies in parental investment, as well as in the recruitment of helping individuals (***McNamara et al., 1999***; ***Paquet and Smiseth, 2017***; ***Pilakouta et al., 2016***; ***Rauter and Moore, 2004***). For dominant females, they have no compensation for the loss of mates by adjusting their investment to the current brood as they may already work at a maximum level of care (Creighton et al., 2014; ***Eggert and Müller, 2000***; ***Smiseth et al., 2005***). The fitness costs incurred by recruiting other group members as new breeding partners may outweigh the benefits of having subordinates, because there occurs a severe competition in reproduction between females (***Eggert and Müller, 2000***; ***Paquet and Smiseth, 2017***; ***Ratz and Smiseth, 2018***). In contrast, dominant males may partially compensate for a reduction in the level of care by their dominant partners, because the costs of mate loss may be offset by the benefits of recruiting other females or by increasing the level of care provided by themselves (***McNamara et al., 1999***; ***Rauter & Moore, 2004***; ***Smiseth et al., 2005***; ***Suzuki & Nagano, 2009***; ***Trumbo & Valletta, 2007***).

### Dominant males, but not dominant females, benefit from staying with subordinates in groups

Our results support that dominants could effectively restrict subordinate’s access to the resources, but such restriction is limited and dominants pay costs from living with subordinates in communal breeding (i.e. pay-from-staying; ***Fig. 5 and 6; Table S5 and S6***). In the absence of its dominant partner, a single remaining dominant individual pays increased costs by the presence of subordinates. This may be due to the inability of single dominants to control subordinates (***Eggert and Müller, 1992***; ***Monnin and Peeters, 1999***; ***Shen and Reeve, 2010***), as well as a behavioural difference in response to the loss of mates between sexes (i.e. sex compensation in parental investment; ***Bruintjes et al., 2013***; ***Russell et al., 2008***). Our results highlight the importance of parental investment by dominant females on reproductive benefits of groups, e.g. lighter offspring in the absence of dominant females (Fig. 6; Table S6). However, such costs in reproduction due to the absence of dominant females may not be fully offset by the benefits of increased brood care by dominant males (***Cotter et al., 2011***; ***Lock et al., 2004***; ***Ratz et al., 2020***; ***Trumbo and Valletta, 2007***). By directly assessing the role of subordinate presence on group productivity when one of dominant individuals is absent, we suggest that an enhanced reproductive benefit accrued by the subordinate’s help is more pronounced for dominant males, but not for dominant females (***Fig. 6; Table S7***). The remaining dominant females do not suffer a reduction in reproductive benefits due to the absence of subordinates in communal groups. This indicated that an enhanced effort by dominant females may offset the costs of the absence of their mates and subordinates in group productivity. However, groups with the remaining dominant males and subordinates had enhanced reproductive benefits, i.e. larger brood sizes, compared to groups with the remaining dominant males alone (***Fig. 6; Table S7***). These results show that, in communal groups, the remaining female dominants, but not the remaining male dominants, invest less in parental care and have a worse ability to control subordinates, and then suffer a reduction in reproductive benefits from the presence of subordinates when their mates are absent. Furthermore, dominant males, but not dominant females, may benefit from the help of subordinates, because the dominant males’ costs associated with the loss of dominant mates may be offset by the benefits of recruiting subordinate females that could actively contribute to current broods (***Bruintjes et al., 2013***; ***Maccoll and Hatchwell, 2003***; ***Russell et al., 2008***; ***Trumbo and Valletta, 2007***; ***Zöttl, Fischer, et al., 2013***). This effect may be due to the benefit/cost of communal breeding being different for females and males in burying beetles (***Liu, Chen, et al., 2020***; ***Ma et al., 2022***; ***Richardson and Smiseth, 2020***; ***Scott, 1998***). In some social groups, focal breeders are expected to benefit from the presence of helpers (***Bruintjes et al., 2013***; ***Maccoll and Hatchwell, 2003***; ***Russell et al., 2008***; ***Zöttl, Fischer, et al., 2013***), such as a reduced workload in parental care and an improved group defence (***Hatchwell and Russell, 1996***; ***Kingma et al., 2010***; ***Maccoll and Hatchwell, 2003***; ***Woxvold and Magrath, 2005***). However, there is no firm evidence for dominants receiving direct benefits from the help of subordinates in burying beetles (***Chen et al., 2020***; ***Eggert and Sakaluk, 2000***; ***Komdeur et al., 2013***; ***Scott, 1994***). In contrast, a large number of studies suggest a mutual tolerance interaction in communal groups, wherein the dominant male may benefit from staying with subordinates, whereas dominant females could reduce reproductive costs by rejecting other individuals from the carcass (***Eggert and Müller, 1992***, ***2011***; ***Müller et al., 2007***; ***Richardson and Smiseth, 2020***; ***Scott, 1998***). Overall, we suggest a sex difference in behaviour between dominant females and males, which leads to different fitness benefits directed to each individual during communal breeding, and even sex differences in parental allocation between current and future reproduction.

### Understanding the evolution of social behaviour: the role of a ‘Tug-of-war’ competition

Our results on burying beetles show evidence of a within-group tug-of-war competition over resources and reproduction between dominant and subordinate individuals, which can be supported from two perspectives (***Eggert et al., 2008***; ***Johnstone and Cant, 1999***; ***Liu, Chan, et al., 2020***; ***Shen and Reeve, 2010***). From the subordinate’s perspective, subordinates may pay by helping to remain within groups (i.e. pay to be tolerated in groups), while the payment is not in exchange for a larger reproductive share. Furthermore, the payment correlates with the level of within-group tug-of-war competition, and we have evidence that the within-group tug-of-war competition becomes intense under a high pressure of interspecific competition, e.g. a low tolerance of dominants towards subordinates in the case of beetles. From the dominant’s perspective, dominants pay costs from living with subordinates in the group without dominant control over reproduction, whereas the costs are different for dominant females and males. This may result in allocation differences in parental and alloparental investment by dominants, and sex differences in response to other group members. This would require a trade-off between cooperative investment and the resolution of conflict. In spite of such a tug-of-war competition, we suggest that either party could benefit from a mutual tolerance interaction in such groups (e.g. the suppression of aggressive behaviour, more investment in cooperation; ***Barker et al., 2012***; ***Eggert and Müller, 2011***; ***Liu, Chan, et al., 2020***; ***Reeve et al., 1998***; ***Shen and Reeve, 2010***). Such mutual tolerance evolves as one of proximate features facilitating the evolution of animal sociality when the relatedness among individuals is low or reproductive constraints are high for each individual (***Eggert and Müller, 2011***; ***Grueter et al., 2020***; ***Lhomme and Hines, 2018***; ***Schülke and Ostner, 2021***; ***Shen and Reeve, 2010***). Our study suggests that this could be explained for the following reasons. First, mutual tolerance can be considered as any iterated dyadic interactions, in which the direct reproductive allocation and shared benefits (e.g. cooperative efforts in carcass preparation and group defence in burying beetles) may involve group member control through direct reward (i.e. pay-to-stay), or punishment (i.e. fierce eviction from the group; ***Bray et al., 2016***; ***Cant et al., 2010***; ***Wong and Balshine, 2011***; ***Zöttl, Heg, et al., 2013***). Dominants could potentially profit from the presence of additional individuals via kin-selected and mutualistic benefits (***Clutton-Brock and Parker, 1995***; ***Liu, Chan, et al., 2020***; ***Ratnieks et al., 2006***; ***Sachs et al., 2004***). Such associated benefits may outweigh the costs from having additional individuals due to ecological constraints and the loss of parentage (***Hamilton and Taborsky, 2005***; ***Hatchwell and Komdeur, 2000***; ***Liu, Chan, et al., 2020***). Second, we suggest that such mutual tolerance is an adaptive consequence of a tug-of-war competition, which depends on the balance between the payoffs obtained by negotiation and those obtained by entering into an escalated fight (***Eggert et al., 2008***; ***Heg and Hamilton, 2008***; ***Johnstone and Cant, 1999***; ***Liu, Chen, et al., 2020***). This may have evolved to yield a higher payoff in terms of group productivity, instead of behaving aggressively in such a tug-of-war competition (***Cant and Johnstone, 1999***; ***Shen and Reeve, 2010***). The amount of resource and reproduction that one individual claims could strongly influence the level of cooperation that renders tolerance profitable for other group members. Subsequently, in a biological context, such social negotiation or contract associated with mutual tolerance should be considered as any behavioural processes by which group members can reach a compromise settlement (or stable behavioural equilibrium) without the escalation of within-group conflict (***Cant and Johnstone, 1999***; ***Reeve et al., 1998***; ***Shen and Reeve, 2010***). This may temporarily coerce all members to contribute more to cooperation and overall group productivity, or even sacrifice their own selfish share (***Clutton-Brock, 1998***; ***Reeve et al., 1998***; ***Zöttl, Heg, et al., 2013***). Therefore, selection may favour enhanced mutual tolerance among group members, where individuals can strategically terminate the escalation of within-group conflict based on the optimal trade-off between the division of reproduction and total group productivity (***Cant and Johnstone, 1999***; ***Shen and Reeve, 2010***). We encourage future research to study how ecological constraints and shared benefits jointly influence helping behaviour of individuals in groups, and how reproductive allocation between individuals determine group stability and productivity. This research will contribute to our understanding of the evolution of animal societies, potentially via the mutual tolerance route.

## Materials and methods

### Animal maintenance

We used burying beetles generated from the outbred F_2_ -F_5_ offspring of wild-caught European burying beetles, *N. vespilloides* in Vosbergen, Eelde (53°08’N, 06°35’E), the Netherlands, during spring of 2017 and 2018. Up to six same-sex adult beetles that descended from same broods were housed in transparent clear plastic boxes (15×10×8.5 cm) filled with moist peat and fed with mealworms (*Tenebrio*) twice a week. All adult beetles and larvae were reared at the Animal Facility of University of Groningen, the Netherlands, and were maintained under a 16 h: 8 h light: dark cycle at 20 ± 2°C throughout.

### General procedure

We conducted double-pair treatments to induce communally breeding events, and each breeding event consisted of two pairs, which each consisted of a female and a male, and these two pairs differed in body size (large and small females and males; supplementary materials: Fig. S1; ***Eggert and Müller, 2000***; ***Komdeur et al., 2013***; ***Ma et al., 2022***). In such communally breeding events, two pairs of individuals would jointly engage in carcass preparation and defence, and they reproduce and raise their offspring together in one shared brood by utilizing a carcass (***Eggert and Müller, 1992***; ***Müller et al., 2007***; ***Sun et al., 2014***). Larger individuals are more likely to become dominants through fights and have greater access to the carcass, compared to smaller individuals (i.e. subordinates), because an adult’s body size is a primary factor that determines its competitive ability and associated dominance status (***Eggert and Müller, 2000***; ***Komdeur et al., 2013***; ***Safryn and Scott, 2000***). This scenario enables a binary social interaction (dominant versus subordinate) due to the establishment of a dominance hierarchy. In such groups, males attempt to copulate with all females present on the carcass (i.e. promiscuity between conspecifics), and subordinate females and males begin to act more stealthily and may have a difference in parental investment due to their reproductive strategies (***Eggert and Müller, 1992***; ***Müller et al., 1990***, ***2007***). In the larval period, subordinate females and males are also found to take care of all developing larvae together with dominants on the carcass (e.g. offspring provisioning), occasionally feeding larvae side-by-side (***Müller et al., 2007***; ***Trumbo and Wilson, 1993***). For each breeding event, we provided with a large carcass (a mouse carcass of 25 g) as a breeding resource, because previous studies showed that *N. vespilloides* beetles are more likely to breed communally on carcasses equal or larger than 25 g (***Eggert and Müller, 1992***; ***Komdeur et al., 2013***). Non-sibling virgin adult beetles, aged approx. 2 weeks old at post-eclosion, were randomly selected for use in our experiments. The mean body size of beetles was recorded by measuring pronotum width (accuracy: 0.001 mm) after three replicates (***Beeler et al., 2002***; ***Schrader et al., 2018***). A similarly-sized female and male that descended from different broods were paired (***Pilakouta et al., 2015***). Prior to the experiments, each pair was kept separately into a plastic box (15×10×8.5 cm) with 2.0 cm of clean peat for 4 – 5 h to ascertain that copulation occurred and females had been inseminated, which enables each pair of individuals to reproduce during communal breeding (***Eggert and Müller, 2000***; ***Komdeur et al., 2013***). Then, we put two pairs of beetles together to create a communally-breeding event, where we ensured that two pairs of beetles differed approx. 10% in size because a dominance hierarchy is more likely to be established when the size difference between opponents is greater than 10% (***Komdeur et al., 2013***; ***Safryn and Scott, 2000***). Large beetles were individually marked by making small holes in the elytra with 00 insect pin. To initiate breeding, we introduced beetles to a breeding box (19×23×12.5 cm) filled with 3.0 – 4.0 mm layer of moist peat and provided with a thawed mouse carcass of 25 ± 1.0 g. Just prior to the experiments, all beetles used were individually weighed (accuracy: 0.1 mg).

### Manipulation experiment: ‘Pay-to-stay’

To test whether subordinates paid to stay by helping dominants in communally breeding groups, we conducted a manipulation experiment, in which we examined the effects of the ‘cooperative behaviour’ status of subordinates (i.e. subordinates assisted dominants in carcass preparation or not) on individual parental behaviour and investment (i.e. time spent on the carcass and weight change) and reproductive success (***see supplementary materials: Fig. S1***). In total, we conducted four types of experimental groups, including three types of treatment groups (dominants with subordinates) in which dominants were assisted by subordinates with either of three ‘cooperative behaviour’ statuses, including (1) ‘non-cooperative’, (2) ‘cooperative’ and (3) ‘pseudo-cooperative’ subordinates, and (4) one control group (dominants without subordinates) where one large pair of individuals bred alone on the carcass. Prior to the experiment, each subordinate pair of individuals had been maintained together to ensure that copulation occurred and females had been inseminated as described in the general procedure above. In each experimental group with ‘cooperative’ subordinates that assisted dominants with carcass burial (n = 28), one large pair of beetles and one small pair of beetles were transferred to a breeding box at the same time to induce a breeding event (***Eggert and Müller, 2011***; ***Komdeur et al., 2013***). For the treatment groups consisting of ‘non-cooperative’ subordinates that did not assist dominants with carcass burial (n = 27), one large pair of beetles was first transferred to a breeding box and given a carcass to initiate breeding at the onset of the experiment, and then one small pair of beetles that were not given a carcass was introduced to a large pair at 36.0 ± 1.0 h after egg-laying (the peak of egg-laying) (***Eggert and Müller, 2000***, ***2011***). To clarify whether it is the subordinates’ assistance with dominants rather than subordinates’ own effort in carcass burial that influenced their subsequent parental investment (e.g. parental investment time and weight change), we also created ‘pseudo-cooperative’ subordinates that did not jointly bury the carcass with dominants, but had buried another carcass by themselves. For the treatment groups with ‘pseudo-cooperative’ subordinates (n = 17), one small pair of beetles that were given a mouse carcass of 15.0 ± 1.0 g but did not lay eggs previously was then introduced to a large pair to induce a communally-breeding event. For control groups (n = 15), one large pair of beetles was transferred to a breeding box and provided with a carcass. For each experimental group, a thawed mouse carcass of 25 ± 1.0 g was given to initiate breeding, and each breeding event occurred in a breeding box (19×23×12.5 cm) filled with 3.0 – 4.0 mm layer of moist peat. All behavioural observations and reproductive measures were recorded as described below.

### Manipulation experiment: the occurrence of interspecific competition with blowfly maggots

To understand whether the occurrence of interspecific competition (with blowfly maggots) mediated individual parental investment, as well as social interaction between individuals, in communal groups, we conducted an experiment to manipulate the absence or presence of blowfly maggots (***Fig. 2***). Further, to examine the combined effects of the level of interspecific competition and the ‘cooperative’ status of subordinates on the outcome of ‘pay-to-stay’, we manipulated the ‘cooperative behaviour’ status of subordinates (see above), and allowed all beetles of each group to communally breed on a clean (see above) or a ‘blowfly maggots’ carcass. In total, we conducted four types of experiments in the absence or presence of blowfly maggots, including three types of experiments (dominants with subordinates) in which dominants were assisted by subordinates with either of three ‘cooperative’ statuses (‘non-cooperative’, ‘cooperative’ and ‘pseudo-cooperative’) in communal groups, and one control group (dominants without subordinates; see above). Prior to the experiment, we created the ‘blowfly maggots’ carcass, by making a small hole (1 cm of diameter) on a 25 ± 1.0 g thawed mouse carcass, and then placing 1.5 ± 0.15 g newly-hatched blowfly maggots (*Calliphora*) into the carcass. Then, we created the ‘cooperative’ status of subordinates, and conducted communally breeding events consisting of two pairs of individuals, according to the detailed protocols above. Each experimental group was given a clean or a pre-treated ‘blowfly maggots’ carcass to initiate a breeding event. To ensure that maggots are still present 24h after the experiment causing a continuous pressure of interspecific competition, we also placed 1.5 ± 0.15 g newly-hatched blowfly maggots into the carcass 24 hours after the start of each experiment. All behavioural observations and reproductive measures were recorded as described below.

### Removal experiment: ‘Pay-from-staying’

To study whether dominants had differential responses in parental behaviour and investment to subordinates when their mates were absent, and whether the remaining dominants paid more costs from the presence of subordinates in communal groups, we experimentally removed either a dominant female (n = 19), a dominant male (n = 19), or both dominants (n = 16) from communally breeding groups as described above. To ascertain that each pair could reproduce in communal-breeding events and to reduce the risk of infanticide committed by others, dominant individuals were removed from its breeding group at 36.0 ± 1.0 h after the first laid-eggs were found (***Eggert and Müller, 2011***). To verify whether the presence of a carcass, but not direct exclusion by dominants, influenced subordinate’s subsequent parental behaviour, e.g. access towards the carcass and providing care on the carcass (e.g. offspring provisioning), both dominant individuals were removed from communal breeding groups. As a control, no dominant individuals were removed from groups (n = 18) until larvae dispersed from the carcass. Further, to test whether a single remaining dominant compensated for the loss of its dominant mate by increasing its investment to the current brood in the absence of subordinates, we created two additional treatment groups. In these groups, either the dominant female (n = 17) or the dominant male (n = 20), as well as both subordinates, were removed from their breeding groups at 36.0 ± 1.0 h after the first laid-eggs were found. All behavioural observations and reproductive measures were recorded as described below.

### Behavioural observation and measures of reproductive success

During the entire duration of observations (from the onset of experimentation until larvae dispersal), each beetle’s presence on or around the carcass was checked twice a day (at 10:00 and 17:00). For each check, we had a continuous observation per group lasting 30 seconds so as to ensure whether or not beetles were providing care on the carcass rather than walking occasionally around the carcass without providing care. In burying beetles, parental care provided by females and males always took place on the carcass and was defined as processing the carcass, guarding the carcass and provisioning larvae present on the carcass (***Smiseth et al., 2005***; ***Wang et al., 2021***). For each individual, we measured parental investment time (the amount of time spent on the carcass by individuals) as the proportion of time each individual spent providing care on the carcass (i.e. carcass preparation and offspring provisioning) during the entire observation period. To measure parental investment for dominant and subordinate individuals in such the binary interaction (within-group tug-of-war competition) of communal groups, we calculated the total amount of parenting investment time by pairs of individuals (the proportion of times that either dominant or subordinate individuals spent on the carcass). We calculated the total group investment (the proportion of time that both dominant and subordinate individuals combined spent providing care on the carcass). At larval dispersal from the carcass, brood size of offspring (total number of larvae) and mean larval weight (total weight of larvae/ brood size) were recorded to assess the reproductive success of groups. On the day of larvae dispersal (when larvae dispersed from the carcass), all surviving adult beetles were individually weighed, and individually collected for subsequent experiments. We then calculated weight change during breeding for each individual ([final weight– initial weight]/ initial weight), which can be viewed as an indicator verifying the amount of parental investment for each individual and individual parental care (***Smiseth et al., 2005***; ***Trumbo and Xhihani, 2015***; ***Wang et al., 2021***). After taking these measures, all dispersing larvae of the same brood were transferred into transparent boxes (19×23×12.5 cm) filled with peat for pupation until emerging as adults.

### Parentage analyses

To investigate whether subordinates had enhanced reproductive benefits (i.e. more offspring produced in one shared brood) by helping dominants (i.e. assist with dominants in carcass burial), we collected adult parents and newly-emerged offspring for parentage analyses. These beetles were from two different types of communal groups with ‘cooperative’ or ‘non-cooperative’ subordinates present (details described above). For both adults and offspring, total genomic DNA was extracted from individual beetle thorax tissue using the method described in ***Richardson et al. (2001)***. These DNA samples were subsequently genotyped for 5 microsatellite loci (Nvesp_A, F, J, M, & P) previously developed for the burying beetle, *N. vespilloides* (***Pascoal and Kilner, 2017***). The microsatellite loci were amplified in one multiplex PCR using a Qiagen multiplex PCR kit and manufacturer’s protocol. Allele sizes were visualised and scored using an AB3730 DNA analyser and Genemapper 4.0 software (Applied Biosystems). Parent-offspring relatedness was assigned by comparing adult and offspring genotypes for the 5 microsatellite loci within communal groups. An adult was excluded as genetic parent if it’s genotyped mismatched with the offspring’s genotype at 1 or more loci. Locus Nvesp_P carried an 0-allele, and this was taken into account when comparing parent-offspring genotypes. In total, 384 offspring from 19 broods were assigned a single mother and a single father within communal groups.

### Statistical analyses

All statistical analyses were conducted using R version 3.6.3 (R Development Core Team 2018). The statistics of the terms that are in the best models with the lowest AIC values are shown using AICstep model selection.

### Subordinates pay to stay with groups in the absence or presence of blowfly maggots

To examine the combined effect of the level of cooperative status of subordinates and interspecific competition (with blowfly maggots) on the parental investment of dominant and subordinate individuals, we used generalized linear models (GLM with a binomial error structure) and linear models (LM) to analyse parental investment time by dominants and subordinates, as well as weight change of individuals, using blowfly maggots presence (yes vs. no), dominance status (dominant vs. subordinate), sex, cooperative status of subordinates (non-cooperative vs. cooperative vs. pseudo-cooperative), brood size of offspring and their interactions as explanatory factors. Also, we used GLM (with a binomial error structure) to analyse the total group investment (i.e. total group investment time) using the cooperative status of subordinates, brood size and their interaction as explanatory factors. Further, to examine whether cooperative behaviour by subordinates (i.e. subordinates assisted dominants with carcass burial) have a generalized effect on individual parental investment during communal breeding, we redefined the ‘helping behaviour’ of subordinates (whether or not subordinates helped in carcass burial), and ’helping behaviour’ to dominants (whether or not dominants received help in carcass burial from others) (***see supplementary materials: Fig. S5; Table S11 and S12***). We ran GLMs or LMs to analyse the reproductive success of groups (i.e. brood size and mean larval weight) and offspring survival at the post-emerging time, using the cooperative status of subordinates (non-cooperative vs. cooperative vs. pseudo-cooperative), or the helping behaviour to dominants (helped vs. non-helped vs. control) as explanatory factors. To examine the effect of the ‘cooperative’ status of subordinates on individual reproductive success, we used GLM (with a binomial error structure) to analyse the percentage of offspring that was produced by individuals using dominance (dominant vs. subordinate), sex, the cooperative status of subordinates (non-cooperative vs. cooperative) and their interactions as explanatory factors. We analysed the number of offspring produced by individuals using LM using dominance, individual time spent providing care and their interactions as explanatory factors.

### Dominants pay from having subordinates

To examine the effect of the removals of dominant females or males on subordinate’s parenting performance, we used GLM (with a binomial error structure) to analyse the amount of care provided by subordinate individuals using removal treatment (removal of dominant female, dominant male, both dominants, and no removals), sex, brood size, and their interactions as explanatory factors. We ran GLM or LM to analyse the amount of time that the remaining dominant individuals spent on the carcass, as well as their weight change before and after parenting, using removal treatment (no removals vs. removal of partner vs. removal of partners and subordinates), sex, brood size, and their interactions as explanatory factors. To study the effect of the removal of dominant females and males on the reproductive success of groups, we ran LMs to analyse brood size and weight of offspring, and mean larval weight, using the removal treatments (no removal, removal of dominant females, removal of dominant males, and removal of both dominants) as an explanatory factor. To examine whether the remaining dominant individuals pay costs or obtain benefits from the presence of subordinates when their dominant partners were absent, we ran LMs to analyse the reproductive success of groups using the removal treatment of dominant males (removal of dominant males, and of dominant males and subordinates), or the removal treatment of dominant females (removal of dominant females, and of dominant males and subordinates) as an explanatory factor.

### Competitive interactions between dominants and subordinates

To investigate whether there occurred a ‘tug-of-war’ competition with respect to parental investment and access towards the carcass between dominant and subordinate individuals, we ran a GLM investigating the relationship of the amount of care provided by subordinates with the amount of time spent by dominants across all treatments. We ran a GLM to analyse the total amount of time spent by communal groups (dominant and subordinate pairs combined). In these two models, the amount of time spent by dominant pairs, by subordinate pairs, the presence of blowfly maggots (yes vs. no) and their interactions were included as explanatory factors. To test whether there occurred a ‘tug-of-war’ competition with respect to weight changes between individuals, we used a LM to analyse individual weight change using dominance (dominant vs. subordinate), the presence of blowfly maggots (yes vs. no), the amount of time spent by individuals, by their partners, and their interactions as explanatory factors. To further study the effect of the amount of time spent by individuals, as well as weight change of individuals on the reproductive outcome of groups (i.e. brood size of offspring and mean larval weight), we ran LMs using the amount of time spent, as well as weight change, of dominants and subordinates as explanatory factors.

To test whether the absence of dominant individuals influenced the intensity of a tug-of-war competition with respect to access to the carcass and weight gains between dominant and subordinate individuals, we used a GLM to analyse the amount of parenting investment time spent by subordinate pairs, using the amount of time spent by dominant pairs, the removal treatment (removal of dominant female, dominant male, both dominants, and no removals), and their interaction as explanatory factors. We used a LM to analyse individual weight change using dominance (dominant vs. subordinate), the removal treatment, the amount of time spent by individuals, by their partners, and their interactions as explanatory factors. Further, to study whether the intensity of a tug-of-war competition was due to the absence of dominant individuals (i.e. the disequilibrium of the control ability between dominants and subordinates due to the absence of dominant individuals), we ran LMs to analyse the reproductive outcome of groups (i.e. brood size of offspring and mean larval weight). In these models, we used the amount of time spent and weight change of dominants and subordinates, and the removal treatment as explanatory factors.

## Data Availability Statement

Data available from the Dryad Digital Repository: https://doi.org/10.5061/dryad.n8pk0p2xj (***Ma et al., 2022***).

## Ethical Note

The research described in this study adheres to the Guidelines for the Use of Animals in Research, University of Groningen. This study was approved by both the University of Groningen and the “Kraus-Groeneveld Stiching”.

## SUPPLEMENTARY MATERIALS

**Fig. S1.**
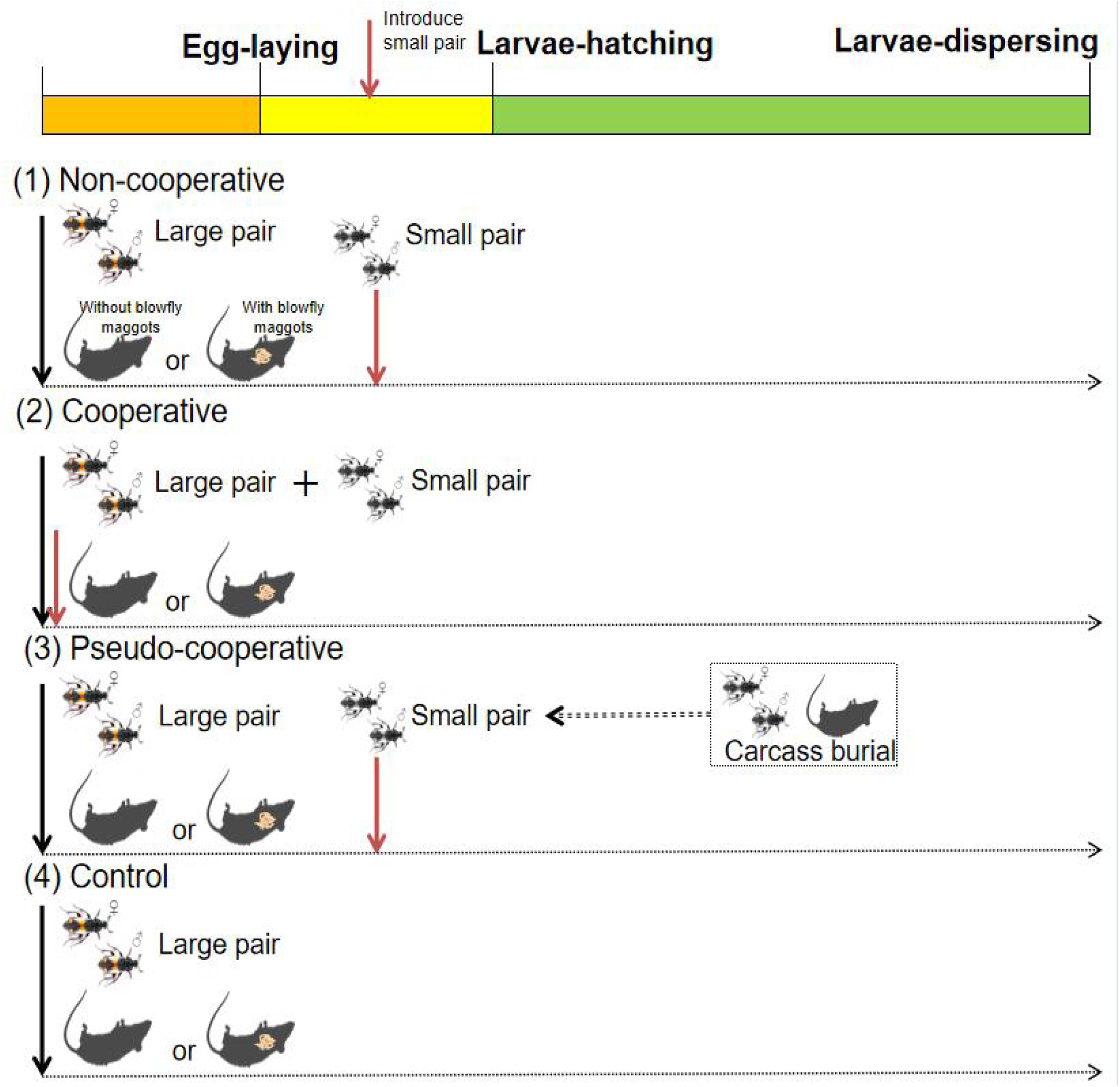
Schematic view of the manipulation experiment in communally breeding burying beetles, *N. vespilloides*. In the manipulation experiment, we conducted a double-pair treatment to induce a communally breeding event, which each consisted of one large pair (dominants) and one small pair (subordinates) of individuals (***Eggert and Müller, 2000***; ***Komdeur et al., 2013***; ***Richardson and Smiseth, 2020***). During carcass preparation, the cooperative status of a small pair of individuals (as subordinates) was manipulated, and the small pair of individuals was assigned to a large pair of individuals (as dominants). The level of interspecific competition (absence or presence of blowfly maggots) was manipulated, and allowed all beetles of each group to communally breed on a clean or a ‘blowfly maggots’ carcass. In all treatments, each small pair of beetles was manipulated and then introduced to a large pair at 36 ± 1h after egg-laying. In total, four types of experimental groups were conducted, including three types of treatment groups in which dominants were assisted by subordinates with either of three ‘cooperative behaviour’ statuses: (1) ‘non-cooperative’ subordinates that were mated and did not assist dominants with carcass burial; (2) ‘cooperative’ subordinates that assisted dominants with carcass burial; and (3) ‘pseudo-cooperative’ subordinates that did not jointly bury the carcass with dominants, but had buried another carcass by themselves. The group without a small pair of subordinates was conducted as control with one large pair of beetles bred alone on the carcass.

**Fig. S2.**
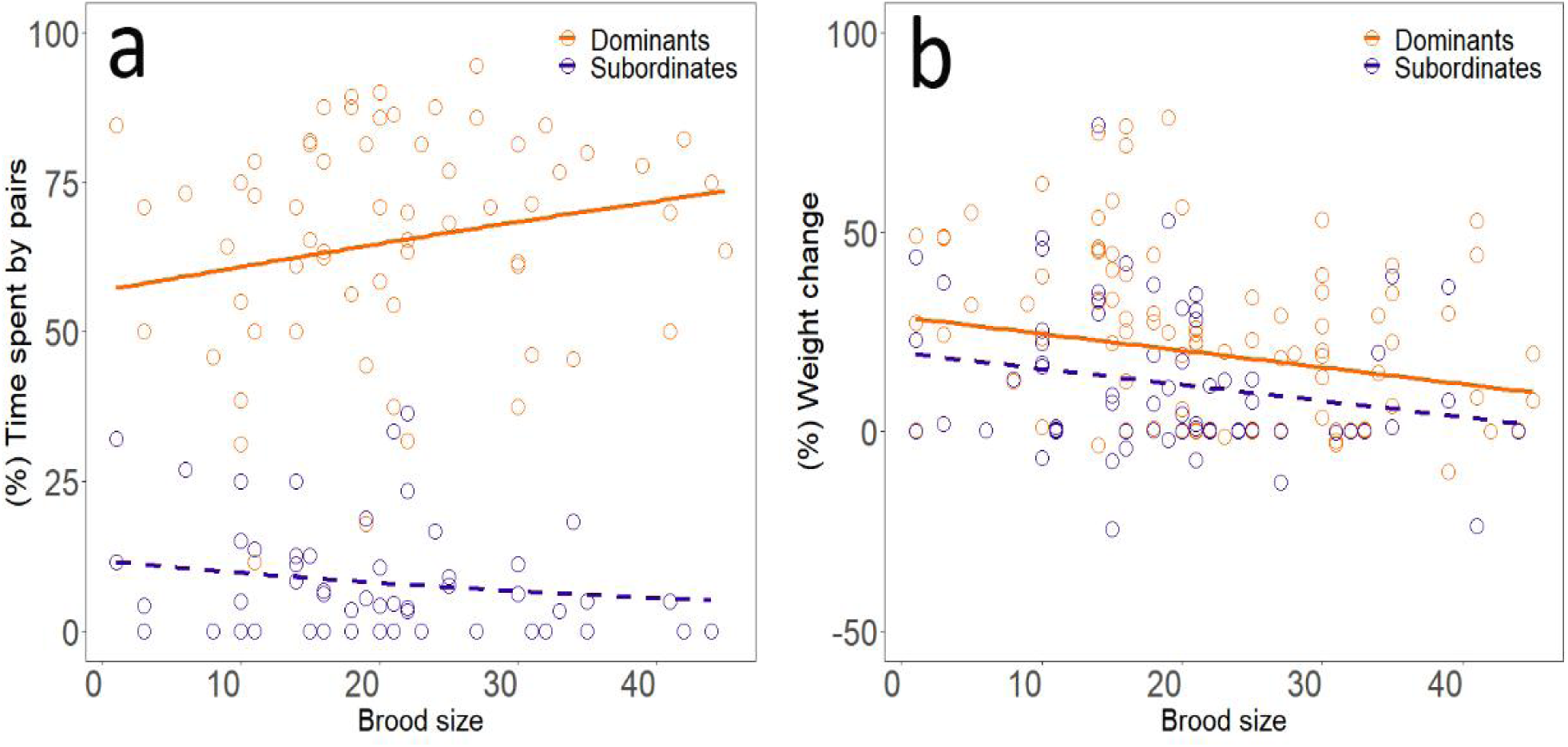
Relationships of (a) time spent on the carcass and (b) weight change of individuals with the brood size of offspring. Dominant individuals (orange lines and dots) and subordinate individuals (purple lines and dots). Solid lines indicate significant correlations, and dotted lines are non-significant correlations. See Table S1 for more statistical details.

**Fig. S3.**
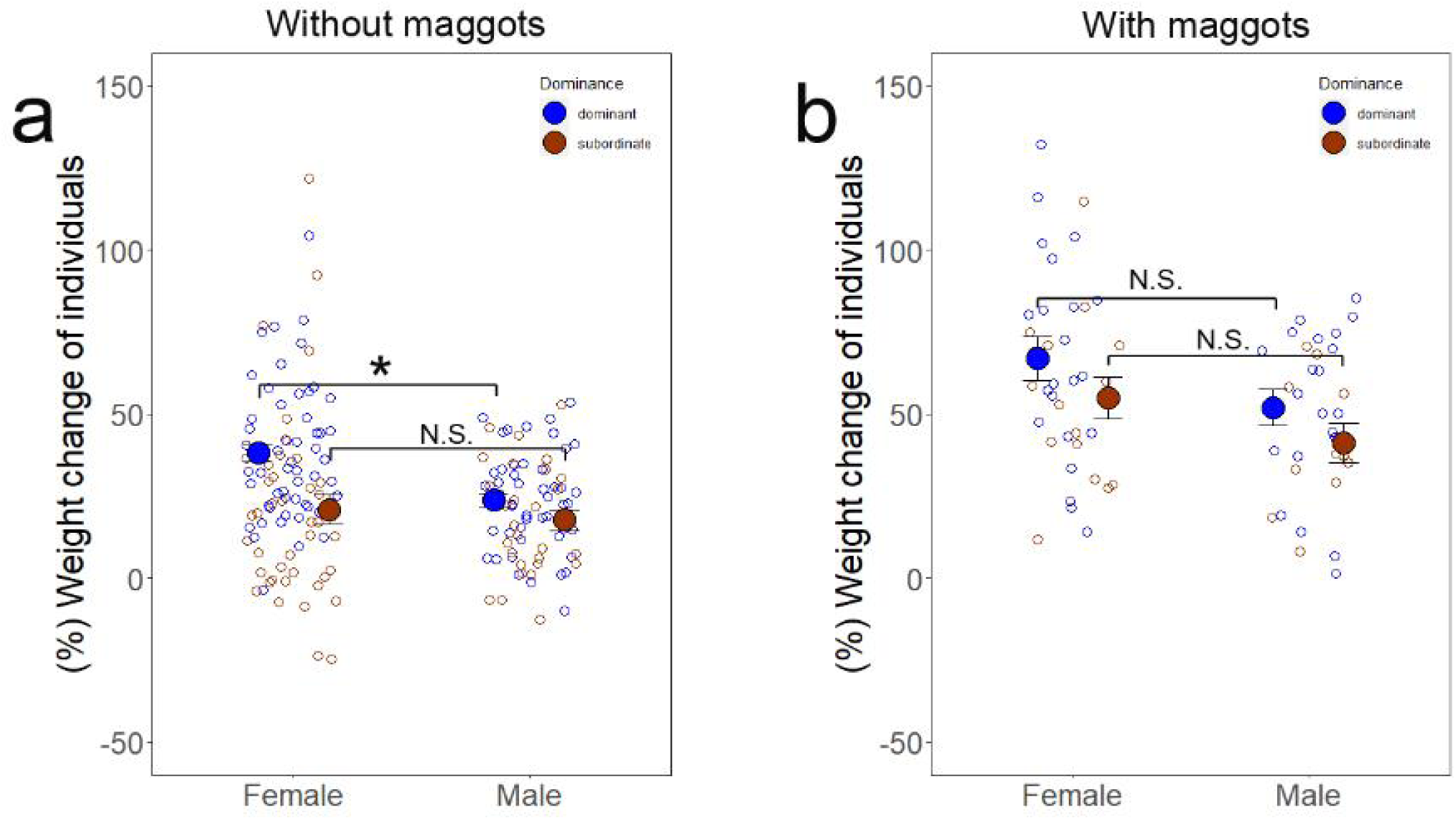
Weight change (%) of dominant and subordinate females and males in (a) the absence (without maggots) and (b) presence of blowfly maggots (with maggots) during breeding. X-axis labels indicate female and male individuals. Dominance status: dominant (blue dots) and subordinate individuals (red dots). Notes: Asterisks above error bar indicate significance: *, p < 0.05; **, p < 0.01; ***, p < 0.001. Raw data and means ± standard errors (SE) are shown. See Table S1 for more statistical details.

**Fig. S4.**
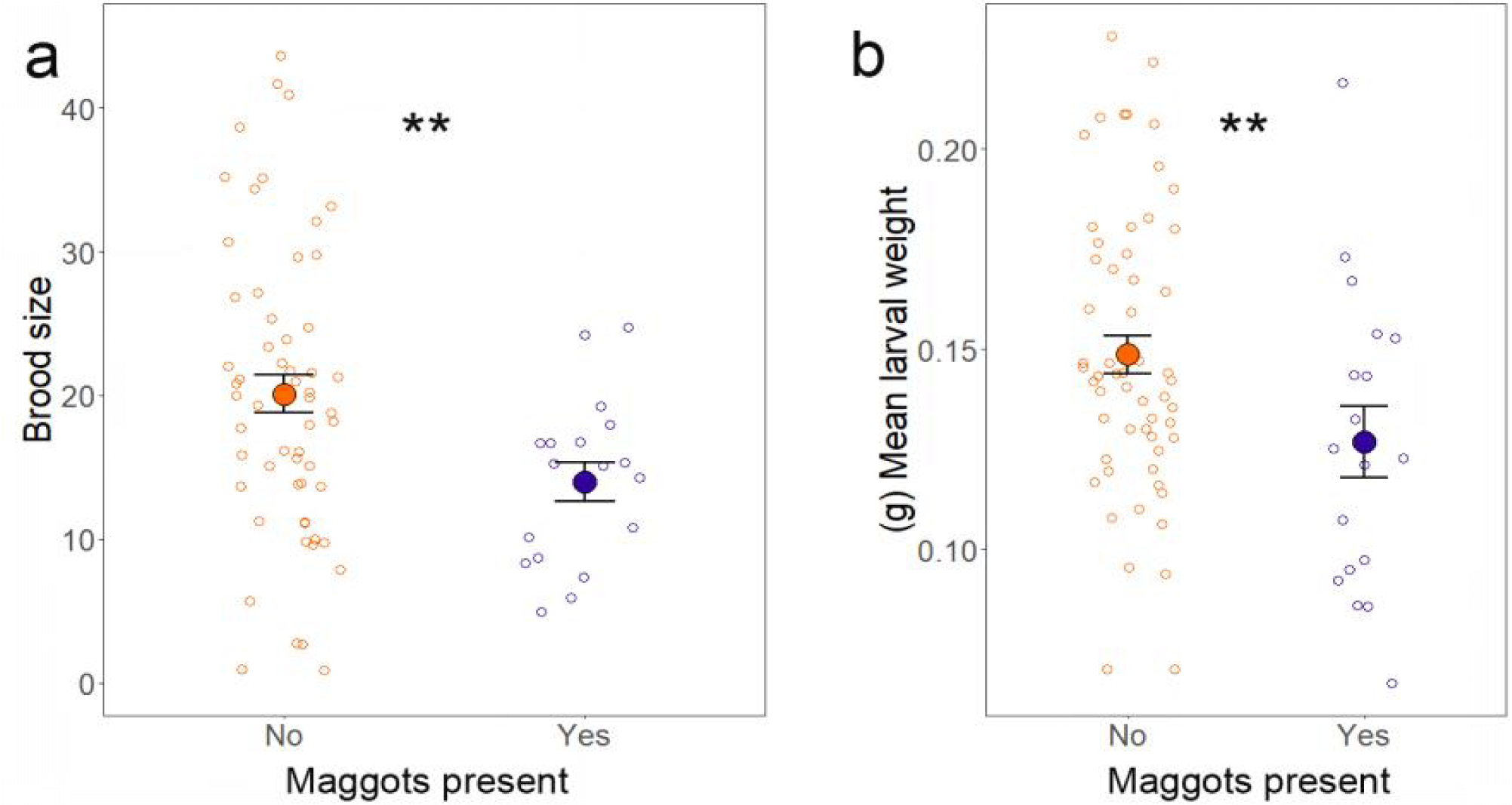
Effects of the presence of blowfly maggots on (a) brood size and (b) mean larval weight of groups in communal breeding. X-axis labels indicate the absence (maggots present: No; orange dotes) and presence (maggots present: Yes; purple dots) of blowfly maggots. Notes: Asterisks above error bar indicate significance: *, p < 0.05; **, p < 0.01; ***, p < 0.001. Raw data and means ± standard errors (SE) are shown. See Table S2 for more statistical details.

**Table S1.**
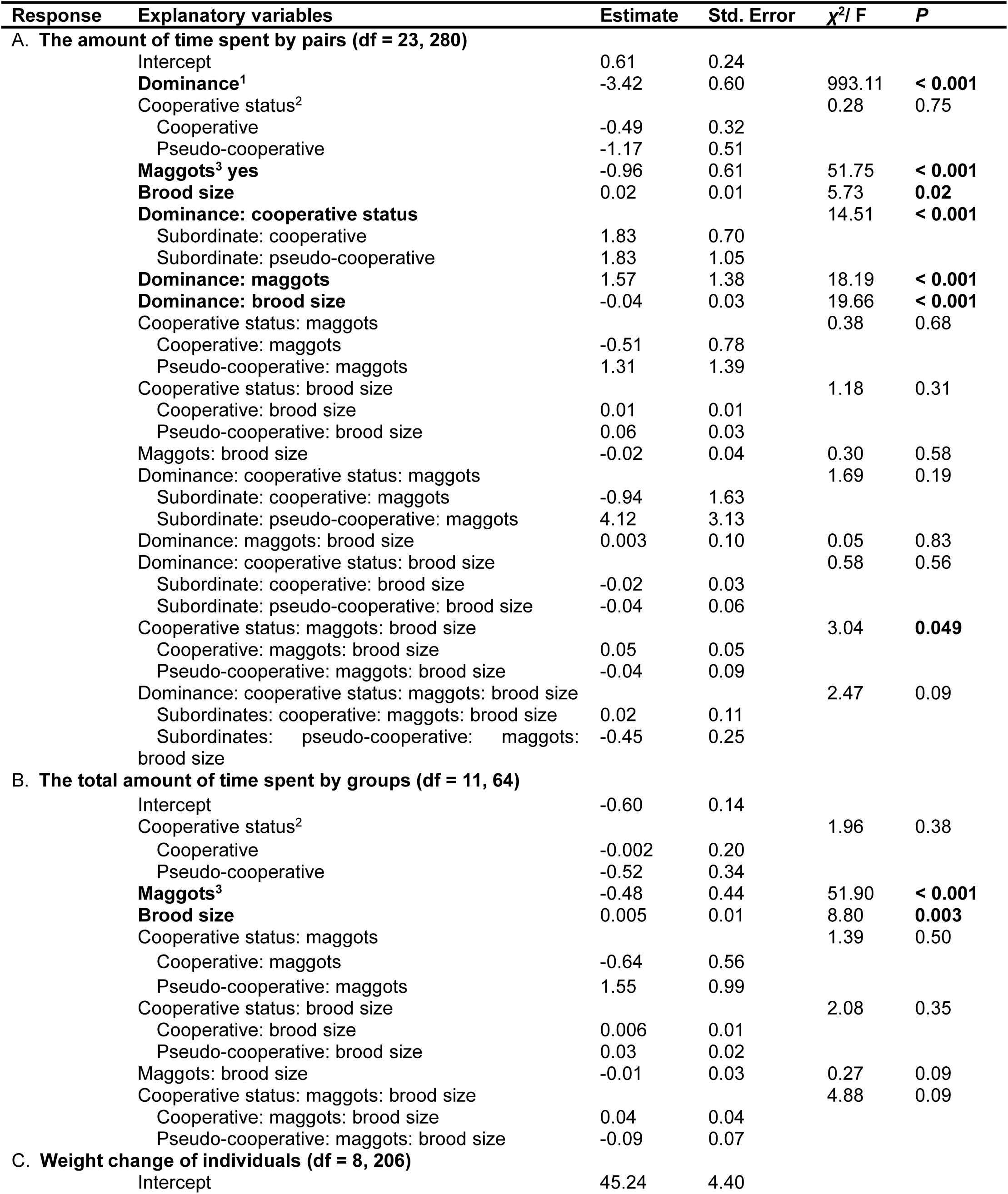
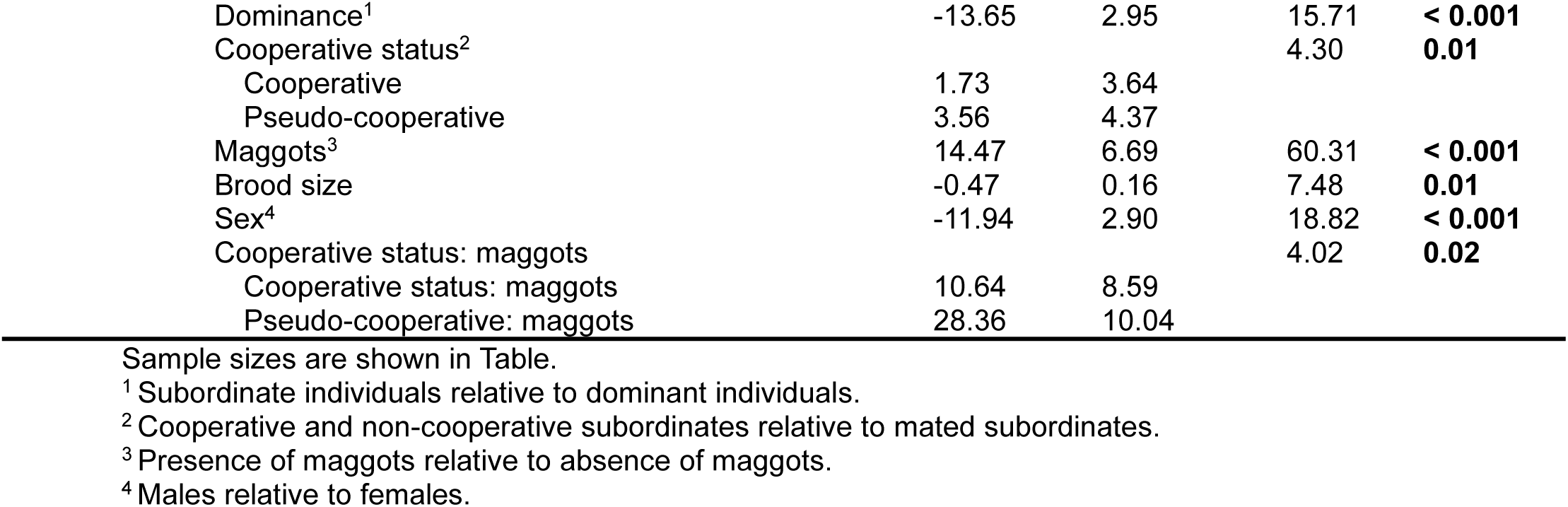
The effects of dominance, the cooperative status of subordinates, sex and brood size of offspring on (A) the amount of time spent by dominants and subordinates, (B) the total amount of time spent by groups, and (C) the weight change of individuals in communally breeding burying beetles, N. vespilloides.

| Response | Explanatory variables | Estimate | Std. Error | $\chi^2/ F$ | P |
| --- | --- | --- | --- | --- | --- |
| **A. The amount of time spent by pairs (df = 23, 280)** |  |  |  |  |  |
|  | Intercept | 0.61 | 0.24 |  |  |
|  | **Dominance<sup>1</sup>** | -3.42 | 0.60 | 993.11 | **< 0.001** |
|  | Cooperative status<sup>2</sup> |  |  | 0.28 | 0.75 |
|  | Cooperative | -0.49 | 0.32 |  |  |
|  | Pseudo-cooperative | -1.17 | 0.51 |  |  |
|  | **Maggots<sup>3</sup> yes** | -0.96 | 0.61 | 51.75 | **< 0.001** |
|  | **Brood size** | 0.02 | 0.01 | 5.73 | **0.02** |
|  | **Dominance: cooperative status** |  |  | 14.51 | **< 0.001** |
|  | Subordinate: cooperative | 1.83 | 0.70 |  |  |
|  | Subordinate: pseudo-cooperative | 1.83 | 1.05 |  |  |
|  | **Dominance: maggots** | 1.57 | 1.38 | 18.19 | **< 0.001** |
|  | **Dominance: brood size** | -0.04 | 0.03 | 19.66 | **< 0.001** |
|  | Cooperative status: maggots |  |  | 0.38 | 0.68 |
|  | Cooperative: maggots | -0.51 | 0.78 |  |  |
|  | Pseudo-cooperative: maggots | 1.31 | 1.39 |  |  |
|  | Cooperative status: brood size |  |  | 1.18 | 0.31 |
|  | Cooperative: brood size | 0.01 | 0.01 |  |  |
|  | Pseudo-cooperative: brood size | 0.06 | 0.03 |  |  |
|  | Maggots: brood size | -0.02 | 0.04 | 0.30 | 0.58 |
|  | Dominance: cooperative status: maggots |  |  | 1.69 | 0.19 |
|  | Subordinate: cooperative: maggots | -0.94 | 1.63 |  |  |
|  | Subordinate: pseudo-cooperative: maggots | 4.12 | 3.13 |  |  |
|  | Dominance: maggots: brood size | 0.003 | 0.10 | 0.05 | 0.83 |
|  | Dominance: cooperative status: brood size |  |  | 0.58 | 0.56 |
|  | Subordinate: cooperative: brood size | -0.02 | 0.03 |  |  |
|  | Subordinate: pseudo-cooperative: brood size | -0.04 | 0.06 |  |  |
|  | Cooperative status: maggots: brood size |  |  | 3.04 | **0.049** |
|  | Cooperative: maggots: brood size | 0.05 | 0.05 |  |  |
|  | Pseudo-cooperative: maggots: brood size | -0.04 | 0.09 |  |  |
|  | Dominance: cooperative status: maggots: brood size |  |  | 2.47 | 0.09 |
|  | Subordinates: cooperative: maggots: brood size | 0.02 | 0.11 |  |  |
|  | Subordinates: pseudo-cooperative: maggots: brood size | -0.45 | 0.25 |  |  |
| **B. The total amount of time spent by groups (df = 11, 64)** |  |  |  |  |  |
|  | Intercept | -0.60 | 0.14 |  |  |
|  | Cooperative status<sup>2</sup> |  |  | 1.96 | 0.38 |
|  | Cooperative | -0.002 | 0.20 |  |  |
|  | Pseudo-cooperative | -0.52 | 0.34 |  |  |
|  | **Maggots<sup>3</sup>** | -0.48 | 0.44 | 51.90 | **< 0.001** |
|  | **Brood size** | 0.005 | 0.01 | 8.80 | **0.003** |
|  | Cooperative status: maggots |  |  | 1.39 | 0.50 |
|  | Cooperative: maggots | -0.64 | 0.56 |  |  |
|  | Pseudo-cooperative: maggots | 1.55 | 0.99 |  |  |
|  | Cooperative status: brood size |  |  | 2.08 | 0.35 |
|  | Cooperative: brood size | 0.006 | 0.01 |  |  |
|  | Pseudo-cooperative: brood size | 0.03 | 0.02 |  |  |
|  | Maggots: brood size | -0.01 | 0.03 | 0.27 | 0.09 |
|  | Cooperative status: maggots: brood size |  |  | 4.88 | 0.09 |
|  | Cooperative: maggots: brood size | 0.04 | 0.04 |  |  |
|  | Pseudo-cooperative: maggots: brood size | -0.09 | 0.07 |  |  |
| **C. Weight change of individuals (df = 8, 206)** |  |  |  |  |  |
|  | Intercept | 45.24 | 4.40 |  |  |

|  |  |  |  |  |
| --- | --- | --- | --- | --- |
| Dominance<sup>1</sup> | -13.65 | 2.95 | 15.71 | **< 0.001** |
| Cooperative status<sup>2</sup> |  |  | 4.30 | **0.01** |
| Cooperative | 1.73 | 3.64 |  |  |
| Pseudo-cooperative | 3.56 | 4.37 |  |  |
| Maggots<sup>3</sup> | 14.47 | 6.69 | 60.31 | **< 0.001** |
| Brood size | -0.47 | 0.16 | 7.48 | **0.01** |
| Sex<sup>4</sup> | -11.94 | 2.90 | 18.82 | **< 0.001** |
| Cooperative status: maggots |  |  | 4.02 | **0.02** |
| Cooperative status: maggots | 10.64 | 8.59 |  |  |
| Pseudo-cooperative: maggots | 28.36 | 10.04 |  |  |

Sample sizes are shown in Table.
<sup>1</sup> Subordinate individuals relative to dominant individuals.
<sup>2</sup> Cooperative and non-cooperative subordinates relative to mated subordinates.
<sup>3</sup> Presence of maggots relative to absence of maggots.
<sup>4</sup> Males relative to females.

**Table S2.** Effects of the cooperative status of subordinates (whether subordinates assist with dominants in carcass burial) and the presence of blowfly maggots on (A) the probability of reproductive success, (B) the brood size, (C) the brood weight and (D) the mean larval weight of groups, in communally breeding burying beetles, N. vespilloides.

| Response | Explanatory variables | Estimate | Std. Error | $\chi^2/ F$ | P |
| --- | --- | --- | --- | --- | --- |
| **A. Probability of reproductive success (Yes/No) (df = 5, 97)** |  |  |  |  |  |
|  | Intercept | 1.70 | 0.54 |  |  |
|  | **Maggots<sup>1</sup>** | -1.70 | 0.83 | 8.89 | **0.003** |
|  | Cooperative status<sup>2</sup> |  |  | 0.04 | 0.98 |
|  | Cooperative | -0.47 | 0.69 |  |  |
|  | Non-cooperative | 0.78 | 1.17 |  |  |
|  | Maggots: cooperative status |  |  | 4.03 | 0.13 |
|  | Maggots: cooperative | 1.45 | 1.16 |  |  |
|  | Maggots: non-cooperative | -1.12 | 1.46 |  |  |
| **B. Brood size (df = 5, 70)** |  |  |  |  |  |
|  | Intercept | 19.59 | 1.84 |  |  |
|  | **Maggots<sup>1</sup>** | -6.24 | 4.61 | 5.92 | 0.02 |
|  | Cooperative status<sup>2</sup> |  |  | 1.68 | 0.19 |
|  | Cooperative | 3.11 | 2.74 |  |  |
|  | Non-cooperative | -3.97 | 3.34 |  |  |
|  | Maggots: cooperative status |  |  | 0.70 | 0.50 |
|  | Maggots: cooperative | -2.39 | 5.97 |  |  |
|  | Maggots: non-cooperative | 4.97 | 6.76 |  |  |
| **C. Brood weight (df = 5, 70)** |  |  |  |  |  |
|  | Intercept | 3.00 | 0.28 |  |  |
|  | **Maggots<sup>1</sup>** | -1.27 | 0.66 | 11.28 | 0.001 |
|  | Cooperative status<sup>2</sup> |  |  | 0.72 | 0.49 |
|  | Cooperative | 0.28 | 0.40 |  |  |
|  | Non-cooperative | -0.61 | 0.48 |  |  |
|  | Maggots: cooperative status |  |  | 1.39 | 0.26 |
|  | Maggots: cooperative | -0.44 | 0.86 |  |  |
|  | Maggots: non-cooperative | 1.05 | 0.97 |  |  |
| **D. Mean larval weight (df = 5, 70)** |  |  |  |  |  |
|  | Intercept | 0.15 | 0.01 |  |  |
|  | **Maggots<sup>1</sup>** | -0.01 | 0.02 | 5.21 | 0.02 |
|  | Cooperative status<sup>2</sup> |  |  | 1.28 | 0.28 |
|  | Cooperative | -0.005 | 0.01 |  |  |
|  | Non-cooperative | -0.004 | 0.02 |  |  |
|  | Maggots: cooperative status |  |  | 2.10 | 0.13 |
|  | Maggots: cooperative | -0.03 | 0.02 |  |  |
|  | Maggots: non-cooperative | 0.02 | 0.02 |  |  |

Sample sizes are shown in Table.
<sup>1</sup> Presence of maggots relative to absence of maggots.
<sup>2</sup> Cooperative and non-cooperative subordinates relative to pseudo-cooperative subordinates.

**Table S3.** The effects of dominance, sex, the cooperative status (whether subordinates helped dominants in carcass burial) and the amount of time spent providing care by individuals on (A) the percentage of offspring provided by individuals, (B) number of offspring produced by individuals (irrespective of individual sex), (C) number of offspring produced by pairs and (D) number of offspring produced by individuals, of dominance, sex and the number of offspring produced by individuals on (E) the amount of time spent by individuals, in communally breeding burying beetles, *N. vespilloides*.

| Response | Explanatory variables | Estimate | Std. Error | $\chi^2/ F$ | P |
| --- | --- | --- | --- | --- | --- |
| **A. Percentage of offspring produced by individuals (df = 7, 68)** |  |  |  |  |  |
|  | Intercept | 2.29 | 0.68 |  |  |
|  | Dominance<sup>1</sup> | -4.65 | 0.97 | 1041.27 | < 0.001 |
|  | Sex<sup>2</sup> | < 0.001 | 0.96 | 0.03 | 0.95 |
|  | Cooperative status<sup>3</sup> | -0.97 | 0.81 | 0.13 | 0.89 |
|  | Dominance: sex | -0.16 | 1.40 | 29.95 | 0.04 |
|  | Dominance: cooperative status | 2.01 | 1.16 | 8.46 | 0.26 |
|  | Sex: cooperative status | 1.54 | 1.32 | 0.17 | 0.87 |
|  | Dominance: sex: cooperative status | -2.93 | 1.91 | 16.17 | 0.12 |
| **B. Number of offspring produced by individuals (df = 3, 72)** |  |  |  |  |  |
|  | Intercept | 2.63 | 0.54 |  |  |
|  | Dominance<sup>1</sup> | -1.92 | 0.64 | 524.35 | < 0.001 |
|  | Time spent providing care | 0.003 | 0.01 | 1.59 | 0.60 |
|  | Dominance: time spent providing care | 0.01 | 0.03 | 0.28 | 0.83 |
| **C. Number of offspring produced by pairs (df = 3, 72)** |  |  |  |  |  |
|  | Intercept | 2.33 | 0.54 |  |  |
|  | **Dominance<sup>1</sup>** | -1.15 | 0.59 | 397.85 | < 0.001 |
|  | Time spent providing care | 0.01 | 0.01 | 3.06 | 0.45 |
|  | Dominance: time spent providing care | -0.02 | 0.02 | 2.59 | 0.48 |
| **D. Number of offspring produced by individuals (df = 7, 68)** |  |  |  |  |  |
|  | Intercept | 2.86 | 0.19 |  |  |
|  | Dominance<sup>1</sup> | -2.36 | 0.65 | 89.94 | < 0.001 |
|  | Sex<sup>2</sup> | < 0.001 | 0.27 | < 0.001 | 0.98 |
|  | Cooperative status<sup>3</sup> | -0.09 | 0.27 | 0.14 | 0.71 |
|  | Dominance: sex | -0.14 | 0.95 | 2.75 | 0.10 |
|  | Dominance: cooperative status | 1.03 | 0.77 | 0.87 | 0.35 |
|  | Sex: cooperative status | 0.18 | 0.37 | 0.03 | 0.87 |
|  | Dominance: sex: cooperative status | -1.40 | 1.28 | 1.21 | 0.27 |
| **E. The amount of time spent by individuals (df = 3, 72)** |  |  |  |  |  |
|  | Intercept | 1.38 | 0.30 |  |  |
|  | Dominance<sup>1</sup> | -3.81 | 0.34 | 380.34 | < 0.001 |
|  | Sex<sup>2</sup> | -0.82 | 0.23 | 12.55 | < 0.001 |
|  | Number of offspring provided by individuals | 0.01 | 0.01 | 0.73 | 0.40 |

Sample sizes are shown in Table.
<sup>1</sup> Subordinate individuals relative to dominant individuals.
<sup>2</sup> Males relative to females.
<sup>3</sup> Cooperative subordinate individuals relative to non-cooperative subordinate individuals.

**Table S4.** The removals of dominant females and males influencing (A) the amount of time spent by subordinate individuals, (B) the weight change of subordinates, (C) the amount of time spent by subordinate pairs, (D) the amount of time spent by dominant pairs and (D) the total amount of time spent by groups in communally breeding burying beetles, N. vespilloides.

| Response | Explanatory variables | Estimate | Std. Error | $\chi^2/ F$ | P |
| --- | --- | --- | --- | --- | --- |
| **A. The amount of time spent by subordinate individuals (df = 11, 135)** |  |  |  |  |  |
|  | Intercept | -0.74 | 0.38 |  |  |
|  | **Removal treatment<sup>1</sup>** |  |  | 5.12 | **0.002** |
|  | Dom.female removal | -0.44 | 0.59 |  |  |
|  | Dom.male removal | -2.79 | 0.91 |  |  |
|  | No removal | -0.80 | 0.59 |  |  |
|  | Sex<sup>2</sup> | -0.75 | 0.50 | 1.53 | 0.22 |
|  | Brood size | 0.05 | 0.03 | 0.83 | 0.36 |
|  | Removal treatment: sex |  |  | 1.56 | 0.20 |
|  | Dom.female removal: sex | 0.11 | 0.79 |  |  |
|  | Dom.male removal: sex | 1.66 | 0.83 |  |  |
|  | No removal: sex | 0.21 | 0.76 |  |  |
|  | Removal treatment: brood size |  |  | 2.60 | 0.06 |
|  | Dom.female removal: brood size | -0.01 | 0.05 |  |  |
|  | Dom.male removal: brood size | 0.01 | 0.04 |  |  |
|  | No removal: brood size | -0.06 | 0.04 |  |  |
| **B. Weight change of subordinates (df = 2, 53)** |  |  |  |  |  |
|  | Intercept | 48.84 | 8.41 |  |  |
|  | Sex<sup>2</sup> | -13.00 | 7.41 | 3.31 | 0.07 |
|  | **Brood size** | -0.89 | 0.37 | 5.74 | **0.02** |
| **C. The amount of time spent by subordinate pairs (df = 7, 39)** |  |  |  |  |  |
|  | Intercept | -1.56 | 1.22 |  |  |
|  | **Removal treatment<sup>1</sup>** |  |  | 53.46 | **0.01** |
|  | Dom.female removal | -0.36 | 1.62 |  |  |
|  | Dom.male removal | -1.43 | 1.52 |  |  |
|  | No removal | -0.32 | 1.41 |  |  |
|  | Brood size | 0.08 | 0.08 | 3.08 | 0.41 |
|  | Removal treatment: brood size |  |  | 11.90 | 0.45 |
|  | Dom.female removal: brood size | -0.09 | 0.10 |  |  |
|  | Dom.male removal: brood size | -0.02 | 0.09 |  |  |
|  | No removal: brood size | -0.08 | 0.08 |  |  |
| **D. The amount of time spent by dominant pairs (df = 5, 34)** |  |  |  |  |  |
|  | Intercept | -1.36 | 0.56 |  |  |
|  | **Removal treatment<sup>3</sup>** |  |  | 46.58 | **< 0.001** |
|  | Dom.male removal | 1.90 | 0.75 |  |  |
|  | No removal | 1.18 | 0.66 |  |  |
|  | Brood size | 0.04 | 0.04 | 3.02 | 0.27 |
|  | Removal treatment: brood size |  |  | 14.64 | **0.06** |
|  | Dom.male removal: brood size | -0.08 | 0.04 |  |  |
|  | No removal: brood size | -0.01 | 0.04 |  |  |
| **E. The total amount of time spent by groups (df = 4, 42)** |  |  |  |  |  |
|  | Intercept | -1.56 | 0.18 |  |  |
|  | **Removal treatment<sup>1</sup>** |  |  | 41.82 | **< 0.001** |
|  | Dom.female removal | 0.01 | 0.20 |  |  |
|  | Dom.male removal | 0.49 | 0.18 |  |  |
|  | No removal | 0.73 | 0.18 |  |  |
|  | Brood size | 0.01 | 0.005 | 4.70 | **0.03** |

Sample sizes are shown in Table.
<sup>1</sup> Removals of dominant females and males, and no removal of dominants relative to removals of both dominants.
<sup>2</sup> Males relative to females.
<sup>3</sup> Removal of dominant males, and no removal of dominants relative to removals of dominant females.

**Table S5.** The removals of dominant partners and subordinates influencing (A) the amount of time spent by dominant individuals, (B) the amount of time spent by dominant pairs, (C) the total amount of time spent by groups and (D) the weight changes of dominant individuals in communally breeding burying beetles, *N. vespilloides*.

| Response | Explanatory variables | Estimate | Std. Error | $\chi^2/ F$ | P |
| --- | --- | --- | --- | --- | --- |
| **A. The amount of time spent by dominant individuals (df = 11, 103)** |  |  |  |  |  |
|  | Intercept | -0.88 | 0.36 |  |  |
|  | **Removal treatment<sup>1</sup>** |  |  | 4.23 | **0.02** |
|  | Partner removal | 1.09 | 0.61 |  |  |
|  | Partner-sub removal | 0.85 | 0.68 |  |  |
|  | **Sex<sup>2</sup>** | 0.20 | 0.49 | 8.98 | **0.003** |
|  | **Brood size** | 0.10 | 0.02 | 35.64 | **< 0.001** |
|  | Removal treatment: sex |  |  | 0.69 | 0.50 |
|  | Partner removal: sex | -1.33 | 0.80 |  |  |
|  | Partner-sub removal: sex | -0.51 | 0.89 |  |  |
|  | Removal treatment: brood size |  |  | 0.61 | 0.54 |
|  | Partner removal: brood size | -0.06 | 0.04 |  |  |
|  | Partner-sub removal: brood size | -0.03 | 0.05 |  |  |
|  | Sex: brood size | -0.08 | 0.03 | 2.14 | 0.15 |
|  | Removal treatment: sex: brood size |  |  | 3.82 | **0.02** |
|  | Partner removal: sex: brood size | 0.14 | 0.05 |  |  |
|  | Partner-sub removal: sex: brood size | 0.10 | 0.06 |  |  |
| **B. The amount of time spent by dominant pairs (df = 5, 60)** |  |  |  |  |  |
|  | Intercept | -0.50 | 0.30 |  |  |
|  | **Removal treatment<sup>2</sup>** |  |  | 53.24 | **< 0.001** |
|  | Dom.male+subordinates removal | -0.13 | 0.29 |  |  |
|  | Dom.female removal | -0.49 | 0.31 |  |  |
|  | Dom.male removal | 0.14 | 0.28 |  |  |
|  | No removal | 0.70 | 0.27 |  |  |
|  | Brood size | 0.01 | 0.01 | 1.97 | 0.32 |
| **C. The total amount of time spent by groups (df = 5, 60)** |  |  |  |  |  |
|  | Intercept | -1.48 | 0.17 |  |  |
|  | **Removal treatment<sup>3</sup>** |  |  | 58.92 | **< 0.001** |
|  | Dom.male+subordinates removal | -0.09 | 0.18 |  |  |
|  | Dom.female removal | -0.02 | 0.19 |  |  |
|  | Dom.male removal | 0.47 | 0.16 |  |  |
|  | No removal | 0.72 | 0.16 |  |  |
|  | Brood size | 0.01 | 0.005 | 2.50 | 0.11 |
| **D. Weight change of dominant individuals (df = 8, 62)** |  |  |  |  |  |
|  | Intercept | 53.92 | 65.00 |  |  |
|  | **Removal treatment<sup>1</sup>** |  |  | 1.24 | 0.30 |
|  | Partner removal | -5.44 | 120.08 |  |  |
|  | Partner-sub removal | 140.32 | 100.45 |  |  |
|  | **Sex<sup>2</sup>** | -17.13 | 61.45 | 0.88 | 0.35 |
|  | Brood size | -0.49 | 2.34 | 0.87 | 0.36 |
|  | Removal treatment: sex |  |  | 2.02 | 0.14 |
|  | Partner removal: sex | 4.14 | 90.24 |  |  |
|  | Partner-sub removal: sex | 200.59 | 89.63 |  |  |
|  | Removal treatment: brood size |  |  | 2.02 | 0.14 |
|  | Partner removal: brood size | -0.02 | 5.78 |  |  |
|  | Partner-sub removal: brood size | -9.75 | 4.84 |  |  |

Sample sizes are shown in Table.
<sup>1</sup> Removals of dominant partners, and of dominant partners and subordinates relative to without removals of dominants.
<sup>2</sup> Males relative to females.
<sup>3</sup> Removals of dominant females or males, of dominant males and subordinates, and no removal of dominants relative to removals of dominant females and subordinates.

**Table S6.**
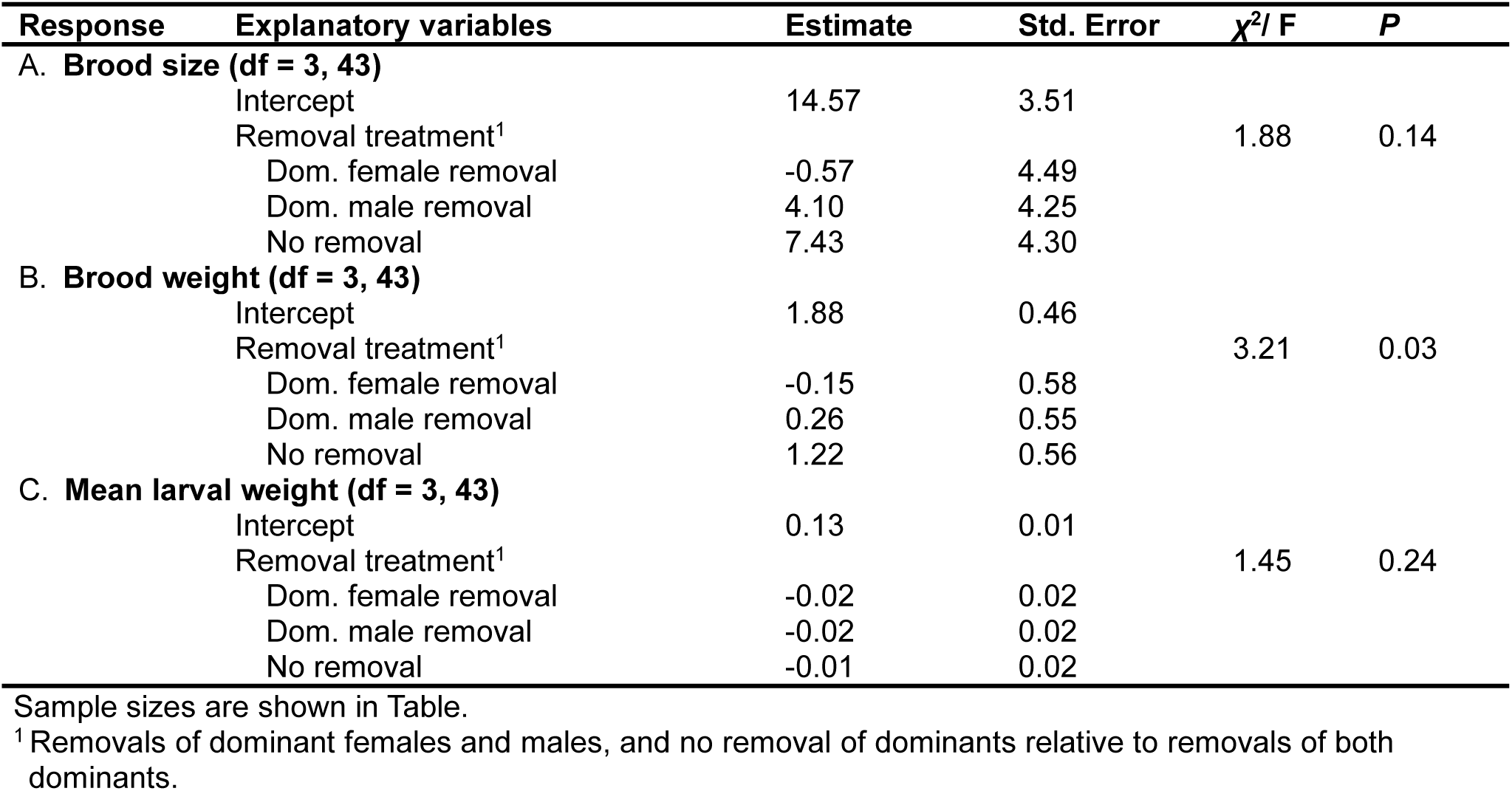
The removals of dominant females and males influencing (A) the brood size, (B) the brood weight and (C) the mean larval weight of groups in communally breeding burying beetles, N. vespilloides.

**Table S7.** The removals of dominant male or female and subordinates influencing (A and D) the brood size, (B and E) the brood weight and (C and F) the mean larval weight of groups in communally breeding burying beetles, *N. vespilloides*.

| Response | Explanatory variables | Estimate | Std. Error | $\chi^2/ F$ | P |
| --- | --- | --- | --- | --- | --- |
| **Do dominant females benefit from the presence of subordinates?** |  |  |  |  |  |
| **A. Brood size (df = 1, 29)** |  |  |  |  |  |
|  | Intercept | 2.73 | 0.11 |  |  |
|  | Removal treatment (dom.male removal)<sup>1</sup> | 0.20 | 0.15 | 1.83 | 0.19 |
| **B. Brood weight (df = 1, 29)** |  |  |  |  |  |
|  | Intercept | 2.11 | 0.22 |  |  |
|  | Removal treatment (dom.male removal)<sup>1</sup> | 0.03 | 0.30 | 0.01 | 0.92 |
| **C. Mean larval weight (df = 1, 29)** |  |  |  |  |  |
|  | Intercept | 0.14 | 0.01 |  |  |
|  | Removal treatment (dom.male removal)<sup>1</sup> | -0.02 | 0.01 | 2.96 | 0.10 |
| **Do dominant males benefit from the presence of subordinates?** |  |  |  |  |  |
| **D. Brood size (df = 1, 20)** |  |  |  |  |  |
|  | Intercept | 21.18 | 2.36 |  |  |
|  | Removal treatment (dom.female removal)<sup>2</sup> | -7.18 | 3.34 | 4.64 | **0.04** |
| **E. Brood weight (df = 1, 20)** |  |  |  |  |  |
|  | Intercept | 2.11 | 0.22 |  |  |
|  | Removal treatment (dom.female removal)<sup>2</sup> | -1.20 | 0.47 | 6.62 | **0.02** |
| **F. Mean larval weight (df = 1, 20)** |  |  |  |  |  |
|  | Intercept | 0.14 | 0.01 |  |  |
|  | Removal treatment (dom.female removal)<sup>2</sup> | -0.02 | 0.01 | 2.60 | 0.12 |

Sample sizes are shown in Table. <sup>1</sup> Removals of dominant males relative to removals of dominant males and subordinates. <sup>2</sup> Removals of dominant females relative to removals of dominant females and subordinates.

**Table S8.** “Tug-of-war” competition affecting (A) the amount of time spent by subordinate pairs, (B) the total amount of time spent by groups and (C and D) the weight change of individuals in communally breeding burying beetles, N. vespilloides.

| Response | Explanatory variables | Estimate | Std. Error | $\chi^2/ F$ | P |
| --- | --- | --- | --- | --- | --- |
| A. The amount of time spent by subordinate pairs (df = 4, 96) |  |  |  |  |  |
|  | Intercept | -2.00 | 0.27 |  |  |
|  | Maggots<sup>1</sup> | -0.56 | 0.20 | 0.62 | 0.43 |
|  | **Cooperative status of subordinates<sup>2</sup>** |  |  | 27.71 | **< 0.001** |
|  | Cooperative | 0.87 | 0.20 |  |  |
|  | Non-cooperative | 0.61 | 0.25 |  |  |
|  | **The amount of time spent by dominant pairs** | -0.02 | 0.004 | 20.24 | **< 0.001** |
| B. The total amount of time spent by groups (df = 5, 95) |  |  |  |  |  |
|  | Intercept | -2.36 | 0.15 |  |  |
|  | Maggots<sup>1</sup> | 0.06 | 0.09 | 74.02 | **< 0.001** |
|  | Cooperative status of subordinates<sup>2</sup> |  |  | 0.10 | 0.95 |
|  | Cooperative | 0.05 | 0.08 |  |  |
|  | Non-cooperative | -0.02 | 0.10 |  |  |
|  | **The amount of time spent by dom. pairs** | 0.02 | 0.002 | 146.74 | **< 0.001** |
|  | **The amount of time spent by sub. pairs** | 0.02 | 0.004 | 36.56 | **< 0.001** |
| C. Weight change of individuals (df = 8, 263) |  |  |  |  |  |
|  | Intercept | 34.57 | 6.81 |  |  |
|  | Dominance<sup>3</sup> | -5.88 | 8.44 | 15.62 | **< 0.001** |
|  | Maggots<sup>1</sup> | 22.82 | 3.89 | 73.12 | **< 0.001** |
|  | Cooperative status of subordinates<sup>2</sup> |  |  | 2.73 | 0.07 |
|  | Cooperative | 1.38 | 3.38 |  |  |
|  | Non-cooperative | 7.87 | 3.90 |  |  |
|  | The amount of time spent by individuals | -7.87 | 8.98 | 0.01 | 0.93 |
|  | The amount of time spent by others | 10.35 | 19.35 | 3.32 | 0.07 |
|  | Dominance: the amount of time spent by individuals | 26.79 | 22.66 | 2.06 | 0.15 |
|  | Dominance: the amount of time spent by others | -29.93 | 21.29 | 1.98 | 0.16 |
| D. Weight change of individuals (df = 7, 264) |  |  |  |  |  |
|  | Intercept | 42.97 | 5.86 |  |  |
|  | Dominance<sup>4</sup> | -11.81 | 2.87 | 15.85 | **< 0.001** |
|  | Maggots<sup>1</sup> | 14.41 | 6.18 | 74.22 | **< 0.001** |
|  | Cooperative status of subordinates<sup>2</sup> |  |  | 2.77 | 0.06 |
|  | Cooperative | 2.48 | 3.61 |  |  |
|  | Non-cooperative | 4.06 | 4.65 |  |  |
|  | **Total time spent by groups** | -33.15 | 14.18 | 6.70 | **0.01** |
|  | Maggots: cooperative status of subordinates |  |  | 1.85 | 0.16 |
|  | Maggots: cooperative | 6.07 | 7.90 |  |  |
|  | Maggots: non-cooperative | 16.00 | 8.40 |  |  |

Sample sizes are shown in Table.
<sup>1</sup> Presence of maggots relative to absence of maggots.
<sup>2</sup> Cooperative and non-cooperative subordinates relative to pseudo-cooperative subordinates.
<sup>3</sup> Dominant individuals relative to subordinate individuals.
<sup>4</sup> Subordinate individuals relative to dominant individuals.

**Table S9.** “Tug-of-war” competition affecting group productivity (i.e. the reproductive success of groups) in communally breeding burying beetles, *N. vespilloides*.

| Response | Explanatory variables | Estimate | Std. Error | $\chi^2/ F$ | P |
| --- | --- | --- | --- | --- | --- |
| **A. Effect of the amount of time spent by pairs on the reproductive success of groups** |  |  |  |  |  |
| Brood size (df = 1, 74) |  |  |  |  |  |
|  | Intercept | 6.72 | 3.31 |  |  |
|  | The amount of time spent by dominant pairs | 0.19 | 0.05 | 14.33 | **< 0.001** |
| Brood weight (df = 1, 74) |  |  |  |  |  |
|  | Intercept | 0.62 | 0.47 |  |  |
|  | The amount of time spent by dominant pairs | 0.03 | 0.01 | 21.54 | **< 0.001** |
| Mean larval weight (df = 1, 74) |  |  |  |  |  |
|  | Intercept | 0.11 | 0.01 |  |  |
|  | The amount of time spent by dominant pairs | < 0.001 | < 0.001 | 8.47 | **0.005** |
| **B. Effect of the total amount of time spent by groups on the reproductive success of groups** |  |  |  |  |  |
| Brood size (df = 1, 74) |  |  |  |  |  |
|  | Intercept | 5.86 | 4.12 |  |  |
|  | The total amount of time spent by groups | 0.37 | 0.11 | 10.29 | **0.002** |
| Brood weight (df = 1, 74) |  |  |  |  |  |
|  | Intercept | 0.39 | 0.59 |  |  |
|  | The total amount of time spent groups | 0.07 | 0.02 | 16.48 | **< 0.001** |
| Mean larval weight (df = 1, 74) |  |  |  |  |  |
|  | Intercept | 0.10 | 0.02 |  |  |
|  | The total amount of time spent by groups | 0.001 | < 0.001 | 7.25 | **0.01** |
| **C. Effect of dominant weight change on the reproductive success of groups** |  |  |  |  |  |
| Brood size (df = 1, 125) |  |  |  |  |  |
|  | Intercept | 23.16 | 1.60 |  |  |
|  | Weight change of dominants | -0.10 | 0.04 | 7.11 | **0.01** |
| Brood weight (df = 1, 125) |  |  |  |  |  |
|  | Intercept | 3.45 | 0.23 |  |  |
|  | Weight change of dominants | -0.02 | 0.005 | 9.24 | **0.002** |
| Mean larval weight (df = 1, 125) |  |  |  |  |  |
|  | Intercept | 0.15 | 0.01 |  |  |
|  | Weight change of dominants | -0.0002 | 0.0001 | 1.11 | 0.29 |
| **D. Effect of subordinate weight change on the reproductive success of groups** |  |  |  |  |  |
| Brood size (df = 1, 86) |  |  |  |  |  |
|  | Intercept | 20.58 | 1.28 |  |  |
|  | Weight change of subordinates | -0.10 | 0.03 | 8.80 | **0.004** |
| Brood weight (df = 1, 86) |  |  |  |  |  |
|  | Intercept | 2.95 | 0.18 |  |  |
|  | Weight change of subordinates | -0.02 | 0.004 | 10.91 | **0.001** |
| Mean larval weight (df = 1, 86) |  |  |  |  |  |
|  | Intercept | 0.14 | 0.005 |  |  |
|  | Weight change of subordinates | -0.0001 | 0.0001 | 0.98 | 0.32 |

Sample sizes are shown in Table.

**Effects of the absence of dominants on the intensity of a tug-of-war competition in communally breeding burying beetles.**

### Statistical analyses

To test whether the absence of dominants influenced the intensity of a tug-of-war competition with respect to access to the carcass and weight gains between dominant and subordinate individuals, we used a GLM to analyse the amount of parenting investment time spent by subordinate pairs, using the amount of time spent by dominant pairs, the removal treatment (removal of dominant female, dominant male, both dominants, and no removals), and their interaction as explanatory factors. We used a LM to analyse individual weight change using dominance, the removal treatment, the amount of time spent by individuals, by their partners, and their interactions as explanatory factors. Further, to study whether the intensity of a tug-of-war competition was due to the absence of dominants (i.e. the disequilibrium of the control ability between dominants and subordinates due to the absence of dominant individuals), we ran LMs to analyse the reproductive success of groups. In these models, we used the amount of time spent and weight change of dominants and subordinates, and the removal treatment as explanatory factors.

**Table S10.** Effect of the loss of dominant individuals on the intensity of a “Tug-of-war” competition over limited resources between dominant and subordinate individuals in communally breeding burying beetles, N. vespilloides.

| Response | Explanatory variables | Estimate | Std. Error | $\chi^2/ F$ | P |
| --- | --- | --- | --- | --- | --- |
| A. The amount of time spent by subordinate pairs (df = 2, 30) |  |  |  |  |  |
|  | Intercept | -0.65 | 0.28 |  |  |
|  | **The amount of time spent by dominant pairs** | -0.05 | 0.01 | 37.55 | **< 0.001** |
|  | **Removal treatment<sup>1</sup>** | 0.62 | 0.26 | 5.81 | **0.02** |
| B. Subordinate weight change (df = 5, 86) |  |  |  |  |  |
|  | Intercept | 35.07 | 9.29 |  |  |
|  | **Sex<sup>2</sup>** | -14.64 | 6.15 | 5.96 | **0.02** |
|  | The amount of time spent by individuals | 0.06 | 0.13 | 0.31 | 0.58 |
|  | Removal treatment<sup>3</sup> |  |  | 0.26 | 0.85 |
|  | Dom.female removal | 4.07 | 9.87 |  |  |
|  | Dom.male removal | 0.69 | 9.56 |  |  |
|  | No removal | -3.67 | 9.66 |  |  |
| C. Dominant weight change (df = 6, 90) |  |  |  |  |  |
|  | Intercept | 77.12 | 71.08 |  |  |
|  | Sex<sup>2</sup> | -5.58 | 47.05 | 0.48 | 0.49 |
|  | The amount of time spent by individuals | 0.58 | 0.48 | 3.07 | 0.08 |
|  | Removal treatment<sup>4</sup> |  |  | 0.75 | 0.56 |
|  | Dom. male + subordinate removal | -81.69 | 64.74 |  |  |
|  | Dom. female removal | -71.87 | 47.77 |  |  |
|  | Dom. male removal | -80.39 | 67.05 |  |  |
|  | No removal | -72.25 | 49.27 |  |  |
| **Effects of the loss of dominant males and parental investment by dominant females on group productivity** |  |  |  |  |  |
| D. Group productivity (brood size) (df = 5, 38) |  |  |  |  |  |
|  | Intercept | 15.68 | 7.42 |  |  |
|  | Dom.female time spent | -0.004 | 0.09 | 1.40 | 0.24 |
|  | Removal treatment<sup>5</sup> |  |  | 2.50 | 0.10 |
|  | Dom.male removal | 9.40 | 9.75 |  |  |
|  | No removal | -19.07 | 10.50 |  |  |
|  | **Dom.female time spent: Removal treatment** |  |  | 6.21 | **0.005** |
|  | Dom.female time spent: Dom.male removal | -0.08 | 0.12 |  |  |
|  | Dom.female time spent: No removal | 0.34 | 0.13 |  |  |
| E. | Group productivity (mean larval weight) (df = 3, 40) |  |  |  |  |
|  | Intercept | 0.10 | 0.02 |  |  |
|  | **Dom.female time spent** | < 0.001 | < 0.001 | 5.52 | **0.02** |
|  | Removal treatment<sup>5</sup> |  |  | 1.74 | 0.19 |
|  | Dom.male removal | -0.02 | 0.01 |  |  |
|  | No removal | 0.003 | 0.01 |  |  |
| F. | Group productivity (brood size) (df = 3, 37) |  |  |  |  |
|  | Intercept | 18.85 | 3.72 |  |  |
|  | Dom.female weight change | -0.09 | 0.07 | 1.24 | 0.27 |
|  | Removal treatment<sup>5</sup> |  |  | 2.56 | 0.09 |
|  | Dom.male removal | 2.37 | 3.31 |  |  |
|  | No removal | 7.24 | 3.25 |  |  |
| G. | Group productivity (mean larval weight) (df = 3, 37) |  |  |  |  |
|  | Intercept | 0.13 | 0.01 |  |  |
|  | Dom.female weight change | < 0.001 | < 0.001 | 1.85 | 0.18 |
|  | Removal treatment<sup>5</sup> |  |  | 1.66 | 0.20 |
|  | Dom.male removal | -0.02 | 0.01 |  |  |
|  | No removal | -0.003 | 0.01 |  |  |
| **Effects of the loss of dominant females and parental investment by dominant males on group productivity** |  |  |  |  |  |
| H. | Group productivity (brood size) (df = 3, 32) |  |  |  |  |
|  | Intercept | 16.37 | 7.60 |  |  |
|  | Dom.male time spent | 0.05 | 0.08 | 0.34 | 0.56 |
|  | Removal treatment<sup>6</sup> |  |  | 2.25 | 0.12 |
|  | Dom.female removal | -5.65 | 4.86 |  |  |
|  | No removal | 3.07 | 5.23 |  |  |
| I. | Group productivity (mean larval weight) (df = 3, 32) |  |  |  |  |
|  | Intercept | 0.14 | 0.03 |  |  |
|  | Dom.male time spent | <-0.001 | < 0.001 | 0.01 | 0.92 |
|  | Removal treatment<sup>6</sup> |  |  | 1.48 | 0.24 |
|  | Dom.female removal | -0.02 | 0.02 |  |  |
|  | No removal | < 0.001 | 0.02 |  |  |
| J. | Group productivity (brood size) (df = 3, 26) |  |  |  |  |
|  | Intercept | 22.59 | 3.51 |  |  |
|  | Dom.male weight change | -0.01 | 0.01 | 0.73 | 0.40 |
|  | Removal treatment<sup>6</sup> |  |  | 2.20 | 0.13 |
|  | Dom.female removal | -7.90 | 4.93 |  |  |
|  | No removal | 1.83 | 4.80 |  |  |
| K. | Group productivity (mean larval weight) (df = 3, 26) |  |  |  |  |
|  | Intercept | 0.14 | 0.11 |  |  |
|  | Dom.male weight change | < -0.001 | < 0.001 | 0.02 | 0.90 |
|  | Removal treatment<sup>6</sup> |  |  | 0.87 | 0.43 |
|  | Dom.female removal | -0.02 | 0.02 |  |  |
|  | No removal | -0.001 | 0.02 |  |  |
| **Effects of the loss of dominants and parental investment by subordinate females on group productivity** |  |  |  |  |  |
| L. | Group productivity (brood size) (df = 4, 42) |  |  |  |  |
|  | Intercept | 12.46 | 4.68 |  |  |
|  | Sub.female time spent | 0.04 | 0.06 | 0.01 | 0.93 |
|  | Removal treatment<sup>3</sup> |  |  | 2.02 | 0.12 |
|  | Dom.female removal | 1.04 | 5.09 |  |  |
|  | Dom.male removal | 5.79 | 4.94 |  |  |
|  | No removal | 8.98 | 4.86 |  |  |
| M. | Group productivity (mean larval weight) (df = 4, 42) |  |  |  |  |
|  | Intercept | 0.14 | 0.02 |  |  |
|  | Sub.female time spent | < -0.001 | < 0.001 | 0.04 | 0.85 |
|  | Removal treatment<sup>3</sup> |  |  | 1.43 | 0.25 |
|  | Dom.female removal | -0.02 | 0.02 |  |  |
|  | Dom.male removal | -0.02 | 0.02 |  |  |
|  | No removal | 0.004 | 0.02 |  |  |
| N. | Group productivity (brood size) (df = 4, 27) |  |  |  |  |
|  | Intercept | 18.46 | 4.50 |  |  |
|  | **Sub.female weight change** | -0.10 | 0.06 | 4.49 | **0.04** |
|  | Removal treatment<sup>3</sup> |  |  | 0.70 | 0.56 |
|  | Dom.female removal | -0.60 | 6.08 |  |  |
|  | Dom.male removal | 4.78 | 4.96 |  |  |
|  | No removal | 5.70 | 5.56 |  |  |
| O. | Group productivity (mean larval weight) (df = 4, 27) |  |  |  |  |
|  | Intercept | 0.14 | 0.02 |  |  |
|  | Sub.female weight change | < -0.001 | < 0.001 | 0.44 | 0.51 |
|  | Removal treatment<sup>3</sup> |  |  | 0.65 | 0.59 |
|  | Dom.female removal | -0.03 | 0.02 |  |  |
|  | Dom.male removal | -0.02 | 0.02 |  |  |
|  | No removal | -0.01 | 0.02 |  |  |
| **Effects of the loss of dominants and parental investment by subordinate males on group productivity** |  |  |  |  |  |
| P. | Group productivity (brood size) (df = 4, 42) |  |  |  |  |
|  | Intercept | 13.69 | 3.91 |  |  |
|  | Sub.male time spent | 0.03 | 0.05 | 0.10 | 0.76 |
|  | Removal treatment<sup>3</sup> |  |  | 1.92 | 0.14 |
|  | Dom.female removal | 0.05 | 4.68 |  |  |
|  | Dom.male removal | 4.40 | 4.33 |  |  |
|  | No removal | 8.02 | 4.48 |  |  |
| Q. | Group productivity (mean larval weight) (df = 4, 42) |  |  |  |  |
|  | Intercept | 0.13 | 0.01 |  |  |
|  | Sub.male time spent | < -0.001 | < 0.001 | 0.10 | 0.76 |
|  | Removal treatment<sup>3</sup> |  |  | 1.42 | 0.25 |
|  | Dom.female removal | -0.02 | 0.02 |  |  |
|  | Dom.male removal | -0.02 | 0.02 |  |  |
|  | No removal | < 0.001 | 0.02 |  |  |
| R. | Group productivity (brood size) (df = 4, 19) |  |  |  |  |
|  | Intercept | 20.63 | 6.70 |  |  |
|  | Sub.male weight change | -0.11 | 0.12 | 0.83 | 0.37 |
|  | Removal treatment<sup>3</sup> |  |  | 0.28 | 0.84 |
|  | Dom.female removal | -3.33 | 7.76 |  |  |
|  | Dom.male removal | 0.90 | 6.82 |  |  |
|  | No removal | 2.37 | 6.98 |  |  |
| S. | Group productivity (mean larval weight) (df = 4, 19) |  |  |  |  |
|  | Intercept | 0.15 | 0.02 |  |  |
|  | Sub.male weight change | < -0.001 | < 0.001 | 0.20 | 0.66 |
|  | Removal treatment<sup>3</sup> |  |  | 1.28 | 0.31 |
|  | Dom.female removal | -0.02 | 0.02 |  |  |
|  | Dom.male removal | -0.02 | 0.02 |  |  |
|  | No removal | -0.03 | 0.02 |  |  |

Sample sizes are shown in Table.
<sup>1</sup> Removal of dominant female relative to removal of dominant male.
<sup>2</sup> Males relative to females.
<sup>3</sup> Removals of dominant females and males, and no removal of dominants relative to removals of both dominants.
<sup>4</sup> Removals of dominant females or males, of dominant males and subordinates, and no removal of dominants relative to removals of dominant females and subordinates.
<sup>5</sup> Removals of dominant males and no removal of dominants relative to removals of dominant males and subordinates.
<sup>6</sup> Removals of dominant females and no removal of dominants relative to removals of dominant females and subordinates.

**Generalized effects of helping behaviour of subordinates on parental investment and the reproductive success of groups**

### Results

We found that the helping behaviour of subordinates in carcass preparation had generalized effects on individual subsequent parental behaviour in dominants and subordinates (***Figure S5; Table S11***), and this was influenced by the pressure of interspecific competition (absence or presence of blowfly maggots; ***supplementary materials: Figure S5; Table S11***). The presence of maggots resulted in a significant reduction in the amount of time spent on the carcass and in a significant increase in weight change for helped (dominants received help in carcass burial from subordinates) and non-helped (dominants did not receive help in carcass burial from subordinates) dominant pairs (z = 3.39, p = 0.002 and z = 3.50, p = 0.001; t = -5.06, p < 0.001 and t = -3.32, p = 0.003; ***Fig. 5a,d; Table S11***), whereas helped and non-helped dominants pairs were similar in the amount of time spent on the carcass and the weight change when maggots were absent or present (time spent on the carcass: z = 1.28, p = 0.65 and z = 1.09, p = 0.77; weight change: t = -0.03, p = 1.00 and t = 0.57, p = 0.97; ***Fig. 5a,d; Table S11***). In the absence of blowfly maggots, dominant pairs that had been helped or not helped spent significantly more time on the carcass (z = -3.28, p = 0.01 and z = -4.31, p < 0.001; ***Fig. S5a; Table S11***), but had no changes in weight (t = -2.14, p = 0.16 and t = -2.13, p = 0.16) compared with control individuals that bred alone (***Fig. S5d; Table S11***). However, this was not the carcass in the presence of blowfly maggots for dominant pairs (z = 0.62, p = 0.96 and z = 1.67, p = 0.38; ***Fig. S5a,d; Table S11***). In the absence of maggots, helpful individuals of subordinates (subordinates helped in carcass burial) spent more time on the carcass compared with non-helpful subordinate pairs (subordinates did not help in carcass burial) (z = -2.28, p = 0.04), while helpful or non-helpful subordinate pairs were similar in the amount of time spent on the carcass when maggots were present (z = -1.18, p = 0.42; ***Fig. S5b; Table S11***). However, the total amount of time spent by groups was similar when dominant pairs had been helped or not (***Fig. S5c; Table S11***). The increasing amount of time spent on the carcass and the fewer gains in weight for dominant pairs, but not for subordinate pairs, resulted in larger broods of offspring in groups (***Table S11***). Groups with subordinate pairs that helped or not in carcass burial were similar in reproductive success of groups in the absence or presence of blowfly maggots (***Table S12***). Groups with helpful subordinates had similar reproductive success of groups in the absence of maggots, but produced significantly lighter larvae in the presence of maggots (t = 2.78, p = 0.04) compared with control individuals that bred alone. However, this was not the case for control groups and groups with non-helpful subordinates (***Table S12***).

#### Statistical analysis

We categorized the ‘helping behaviour’ statuses of subordinates into ‘non-helpful’ (subordinates did not help in carcass burial; ‘non-cooperative’ and ‘pseudo-cooperative’ subordinates) and ‘helpful’ subordinates (subordinates helped in carcass burial previously; ‘cooperative’ subordinates). According to whether dominant individuals received help in carcass burial or not from others, the ‘helping behaviour’ to dominants was categorized into three levels, including ‘helped’ dominants (dominants received help in carcass burial from subordinates; dominants from groups with ‘cooperative’ subordinates), ‘non-helped’ dominants (dominants did not receive help in carcass burial, regardless of whether subordinates helped or not previously; dominants from groups with ‘non-cooperative’ and ‘pseudo-cooperative’ subordinates) and ‘control without subordinates’ dominants (one large pair of individuals, dominants bred alone without subordinates on the carcass). We ran GLMs (with a binomial error structure) to analyse the total amount of time spent by dominant or subordinate pairs. In these analyses, the helping behaviour of subordinates (helpful vs. non-helpful), or the helping behaviour to dominants (helped vs. non-helped vs. control), brood size of offspring and their interactions were included as explanatory factors. We also used LMs to analyse the weight change of dominants and subordinates using sex, the helping behaviour of subordinates (helpful vs. non-helpful), or the helping behaviour to dominants (helped vs. non-helped vs. control), brood size of offspring, and their interactions as explanatory factors.

**Figure S5.**
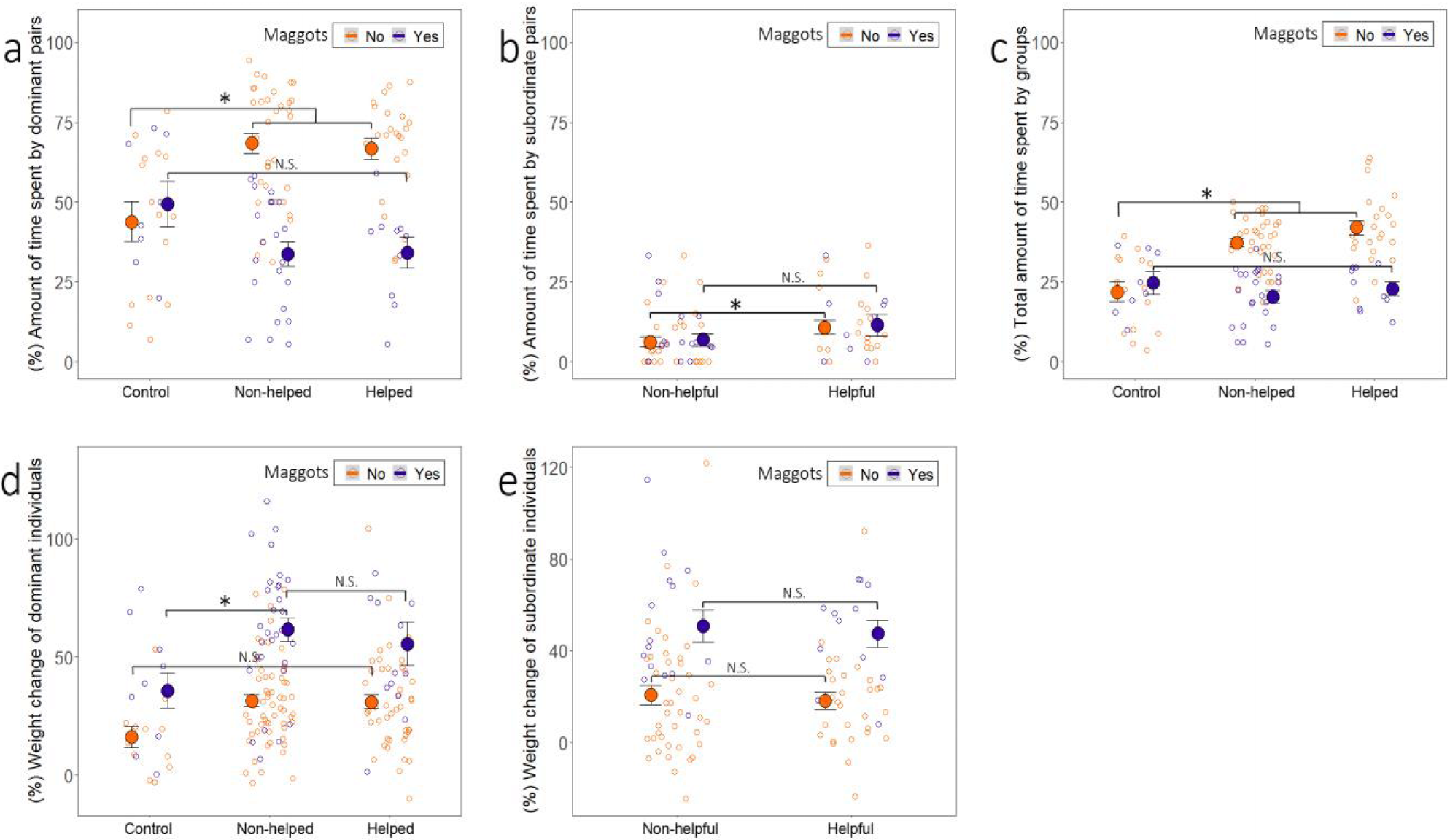
Generalized effects of helping behaviour of subordinates and interspecific competition with blowfly maggots on parental investment in communal breeding of burying beetles, *N. vespilloides*. (a-c) Effects of the level of ‘helping behaviour’ of subordinates (or the level of ‘helping behaviour’ to dominants) on the amount of time spent by dominants (i.e. the dominant pair) **(a)** and subordinates (i.e. the subordinate pair) **(b)**, and on the total amount of time spent by groups **(c)** in the absence and the presence of blowfly maggots. **(d and e)** Effects of the level of ‘helping behaviour’ of subordinates on weight change of dominant (e) and subordinate individuals **(e)**. X-axis labels indicate the level of ‘helping behaviour’ of subordinate individuals: helpful (subordinates help in carcass burial) and non-helpful (subordinates do not help in carcass burial), and the level of ‘helping behaviour’ to dominants: helped (dominants receive help in carcass burial from subordinates), non-helped (dominant do not receive help in carcass burial from subordinates) and control (dominants breed alone without subordinates on the carcass). Blowfly maggots absence (maggots present: No; organe dots) and presence (maggots present: Yes; purple dots), and female (red dots) and male (blue dots) individuals. Notes: Asterisks above error bar indicate significance: *, p < 0.05; **, p < 0.01; ***, p < 0.001. Raw data and means ± standard errors (SE) are shown. See Table S11 for more statistical details.

**Table S11.** The generalized effects of the helping behaviour by subordinates and interspecific competition with blowfly maggots on the amount of time spent provided by dominant (A) and subordinate (B) pairs, (C) the total amount of time spent provided by groups, and the weight change of dominant (D) and subordinate individuals (E) in communally breeding burying beetles, *N. vespilloides*.

| Response | Explanatory variables | Estimate | Std. Error | $\chi^2$/ F | P |
| --- | --- | --- | --- | --- | --- |
| **A. The amount of time spent by dominant pairs (df = 6, 86)** |  |  |  |  |  |
|  | Intercept | -0.73 | 0.29 |  |  |
|  | **Helping behaviour to dominants<sup>1</sup>** |  |  | 4.90 | **0.01** |
|  | Non-helped | 1.08 | 0.25 |  |  |
|  | Helped | 0.83 | 0.25 |  |  |
|  | **Maggots<sup>2</sup>** | 0.38 | 0.32 | 21.50 | **< 0.001** |
|  | **Brood size** | 0.03 | 0.01 | 9.47 | **0.003** |
|  | **Helping behaviour of dominants: maggots** |  |  | 6.69 | **0.002** |
|  | Non-helped: maggots | -1.28 | 0.41 |  |  |
|  | Helped: maggots | -1.40 | 0.42 |  |  |
| **B. The amount of time spent by subordinate pairs (df = 4, 71)** |  |  |  |  |  |
|  | Intercept | -2.32 | 0.38 |  |  |
|  | **Helping behaviour of subordinates<sup>3</sup>** | 0.81 | 0.35 | 5.76 | **0.02** |
|  | Maggots<sup>2</sup> | 0.04 | 0.56 | 0.17 | 0.68 |
|  | Brood size | -0.03 | 0.02 | 3.15 | 0.08 |
|  | Helping behaviour of subordinates: maggots | -0.09 | 0.71 | 0.02 | 0.90 |
| **C. The total amount of time spent by groups (df = 6, 86)** |  |  |  |  |  |
|  | Intercept | -1.44 | 0.17 |  |  |
|  | **Helping behaviour to dominants<sup>3</sup>** |  |  | 13.38 | **< 0.001** |
|  | Non-helpful | 0.72 | 0.15 |  |  |
|  | Helpful | 0.83 | 0.15 |  |  |
|  | **Maggots<sup>2</sup>** | 0.21 | 0.20 | 23.36 | **< 0.001** |
|  | **Brood size** | 0.01 | 0.004 | 6.69 | **0.01** |
|  | **Helping behaviour to dominants: maggots** |  |  | 6.46 | **0.002** |
|  | Non-helped: maggots | -0.70 | 0.25 |  |  |
|  | Helped: maggots | -0.88 | 0.25 |  |  |
| **D. Weight change of dominant individuals (df =7, 141)** |  |  |  |  |  |
|  | Intercept | 30.17 | 7.74 |  |  |
|  | **Sex<sup>3</sup>** | -13.42 | 3.24 | 14.51 | **< 0.001** |
|  | **Helping behaviour to dominants<sup>1</sup>** |  |  | 3.65 | **0.03** |
|  | Non-helped | 13.84 | 6.51 |  |  |
|  | Helped | 13.96 | 6.51 |  |  |
|  | **Maggots<sup>2</sup>** | 19.20 | 8.74 | 50.68 | **< 0.001** |
|  | Brood size | -0.31 | 0.17 | 2.93 | 0.09 |
|  | Helping behaviour of subordinates: maggots |  |  | 0.48 | 0.62 |
|  | Non-helped: maggots | 9.84 | 10.24 |  |  |
|  | Helped: maggots | 4.95 | 11.08 |  |  |
| **E. Weight change of subordinate individuals (df = 5, 82)** |  |  |  |  |  |
|  | Intercept | 34.46 | 6.76 |  |  |
|  | **Sex<sup>4</sup>** | -4.18 | 5.89 | 2.28 | 0.14 |
|  | Helping behaviour of subordinates<sup>3</sup> | 1.05 | 5.43 | 0.07 | 0.79 |
|  | **Maggots<sup>2</sup>** | 34.93 | 8.83 | 19.16 | **< 0.001** |
|  | Brood size | -0.72 | 0.30 | 6.80 | 0.10 |
|  | Sex: maggots | -22.12 | 13.42 | 2.72 | 0.10 |

Sample sizes are shown in Table.
<sup>1</sup> Non-helped and helped dominant individuals relative to control dominant individuals.
<sup>2</sup> Presence of maggots relative to absence of maggots.
<sup>3</sup> Helpful subordinate individuals relative to non-helpful subordinates.
<sup>4</sup> Males relative to females.

**Table S12.**
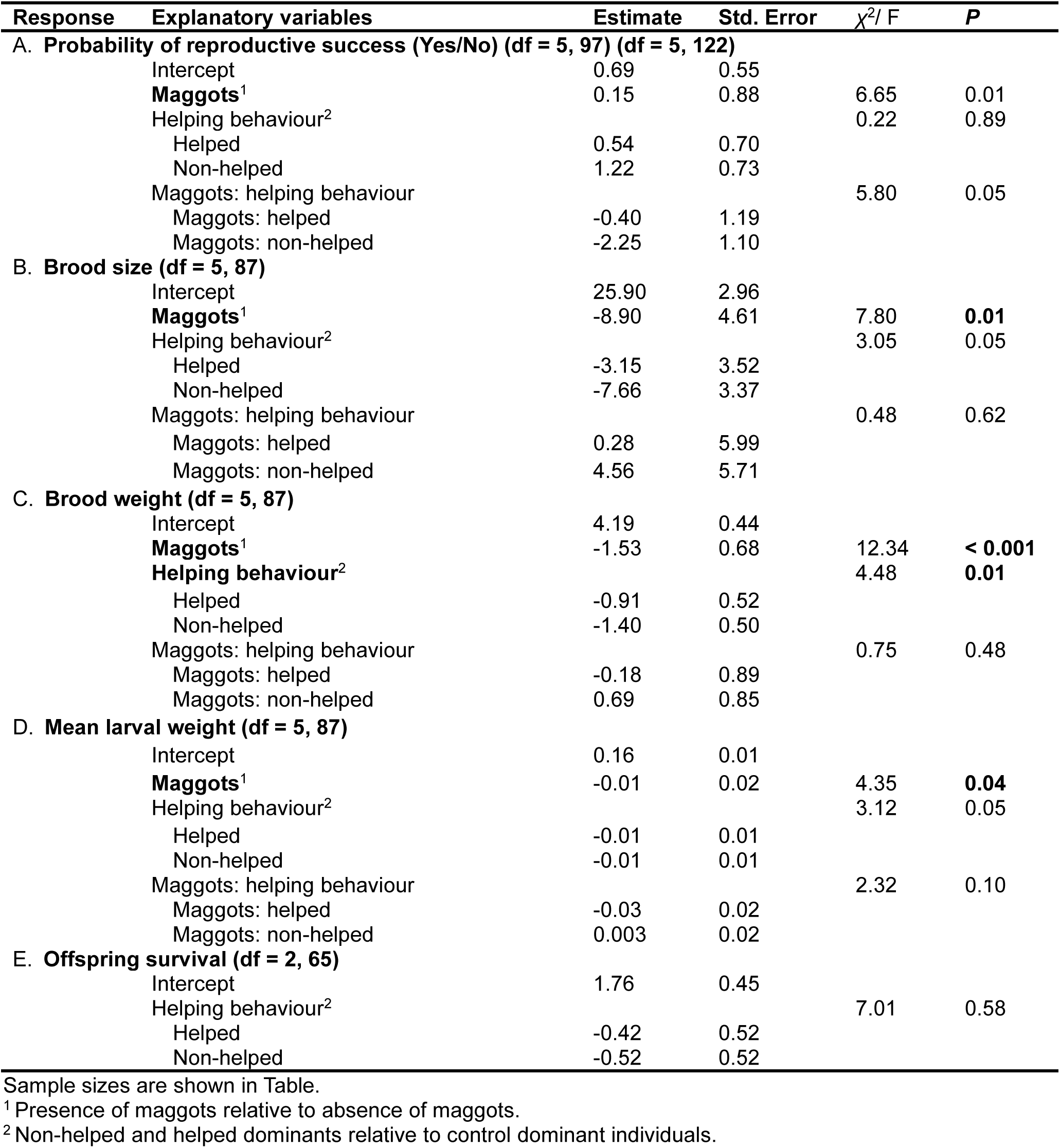
The generalized effects of the helping behaviour dominants (whether dominants receive help from subordinates in carcass burial) and interspecific competition with blowfly maggots on the probability of reproductive success (A), brood size (B), brood weight (C), mean larval weight (D) and offspring survival (E) in communally breeding burying beetles, N. vespilloides.

| Response | Explanatory variables | Estimate | Std. Error | $\chi^2/ F$ | P |
| --- | --- | --- | --- | --- | --- |
| **A. Probability of reproductive success (Yes/No) (df = 5, 97) (df = 5, 122)** |  |  |  |  |  |
|  | Intercept | 0.69 | 0.55 |  |  |
|  | **Maggots**<sup>1</sup> | 0.15 | 0.88 | 6.65 | 0.01 |
|  | Helping behaviour<sup>2</sup> |  |  | 0.22 | 0.89 |
|  | Helped | 0.54 | 0.70 |  |  |
|  | Non-helped | 1.22 | 0.73 |  |  |
|  | Maggots: helping behaviour |  |  | 5.80 | 0.05 |
|  | Maggots: helped | -0.40 | 1.19 |  |  |
|  | Maggots: non-helped | -2.25 | 1.10 |  |  |
| **B. Brood size (df = 5, 87)** |  |  |  |  |  |
|  | Intercept | 25.90 | 2.96 |  |  |
|  | **Maggots**<sup>1</sup> | -8.90 | 4.61 | 7.80 | **0.01** |
|  | Helping behaviour<sup>2</sup> |  |  | 3.05 | 0.05 |
|  | Helped | -3.15 | 3.52 |  |  |
|  | Non-helped | -7.66 | 3.37 |  |  |
|  | Maggots: helping behaviour |  |  | 0.48 | 0.62 |
|  | Maggots: helped | 0.28 | 5.99 |  |  |
|  | Maggots: non-helped | 4.56 | 5.71 |  |  |
| **C. Brood weight (df = 5, 87)** |  |  |  |  |  |
|  | Intercept | 4.19 | 0.44 |  |  |
|  | **Maggots**<sup>1</sup> | -1.53 | 0.68 | 12.34 | **< 0.001** |
|  | Helping behaviour<sup>2</sup> |  |  | 4.48 | **0.01** |
|  | Helped | -0.91 | 0.52 |  |  |
|  | Non-helped | -1.40 | 0.50 |  |  |
|  | Maggots: helping behaviour |  |  | 0.75 | 0.48 |
|  | Maggots: helped | -0.18 | 0.89 |  |  |
|  | Maggots: non-helped | 0.69 | 0.85 |  |  |
| **D. Mean larval weight (df = 5, 87)** |  |  |  |  |  |
|  | Intercept | 0.16 | 0.01 |  |  |
|  | **Maggots**<sup>1</sup> | -0.01 | 0.02 | 4.35 | **0.04** |
|  | Helping behaviour<sup>2</sup> |  |  | 3.12 | 0.05 |
|  | Helped | -0.01 | 0.01 |  |  |
|  | Non-helped | -0.01 | 0.01 |  |  |
|  | Maggots: helping behaviour |  |  | 2.32 | 0.10 |
|  | Maggots: helped | -0.03 | 0.02 |  |  |
|  | Maggots: non-helped | 0.003 | 0.02 |  |  |
| **E. Offspring survival (df = 2, 65)** |  |  |  |  |  |
|  | Intercept | 1.76 | 0.45 |  |  |
|  | Helping behaviour<sup>2</sup> |  |  | 7.01 | 0.58 |
|  | Helped | -0.42 | 0.52 |  |  |
|  | Non-helped | -0.52 | 0.52 |  |  |

Sample sizes are shown in Table.
<sup>1</sup> Presence of maggots relative to absence of maggots.
<sup>2</sup> Non-helped and helped dominants relative to control dominant individuals.

## Notes

### Competing Interest Statement

The authors have declared no competing interest.

